# Long-term expanding human airway organoids for disease modelling

**DOI:** 10.1101/318444

**Authors:** Norman Sachs, Domenique D. Zomer-van Ommen, Angelos Papaspyropoulos, Inha Heo, Lena Böttinger, Dymph Klay, Fleur Weeber, Guizela Huelsz-Prince, Nino Iakobachvili, Marco C. Viveen, Anna Lyubimova, Luc Teeven, Sepideh Derakhshan, Jeroen Korving, Harry Begthel, Kuldeep Kumawat, Emilio Ramos, Matthijs F.M. van Oosterhout, Eduardo P. Olimpio, Joep de Ligt, Krijn K. Dijkstra, Egbert F. Smit, Maarten van der Linden, Emile E. Voest, Coline H.M. van Moorsel, Cornelis K. van der Ent, Edwin Cuppen, Alexander van Oudenaarden, Frank E. Coenjaerts, Linde Meyaard, Louis J. Bont, Peter J. Peters, Sander J. Tans, Jeroen S. van Zon, Sylvia F. Boj, Robert G. Vries, Jeffrey M. Beekman, Hans Clevers

## Abstract

Organoids are self-organizing 3D structures grown from stem cells that recapitulate essential aspects of organ structure and function. Here we describe a method to establish long-term-expanding human airway organoids from broncho-alveolar biopsies or lavage material. The pseudostratified airway organoid epithelium consists of basal cells, functional multi-ciliated cells, mucus-producing goblet cells, and CC10-secreting club cells. Airway organoids derived from cystic fibrosis (CF) patients allow assessment of CFTR function in an organoid swelling assay. Organoid culture conditions also allow gene editing as well as the derivation of various types of lung cancer organoids. Respiratory syncytial virus (RSV) infection recapitulated central disease features and dramatically increases organoid cell motility, found to be driven by the non-structural viral NS2 protein. We conclude that human airway organoids represent versatile models for the in vitro study of hereditary, malignant, and infectious pulmonary disease.

## Introduction

To date, several approaches have been explored to generate mammalian airway organoids^21^. In 1993 Puchelle and colleagues described the first self-organizing 3D structures of adult human airway epithelium in collagen^22^. A first description of the generation of lung organoids from human iPS (induced pluripotent stem) cells was given by Rossant et al. and included the use of *CFTR*-mutant iPS cells as a proof of concept for modeling CF^23^. Snoeck and colleagues designed an improved four-stage protocol^24^, while Spence and colleagues^25^ followed a modified trajectory to generate mature lung organoids, containing basal, ciliated, and club cells. These cultures were stable for up to several months and resembled proximal airways. Hogan and colleagues reported the first adult stem cell-based murine bronchiolar lung organoid culture protocol, involving Matrigel supplemented with EGF^26^. Single basal cells isolated from the trachea grew into *tracheospheres* consisting of a pseudostratified epithelium with basal and ciliated luminal cells. These organoids could be passaged at least twice. No mature club, neuroendocrine, or mucus-producing cells were observed^26^. In a later study, this clonal 3D organoid assay was used to demonstrate that IL-6 treatment resulted in the formation of ciliated cells at the expense of secretory and basal cells^27^. Tschumperlin and colleagues combined human adult primary bronchial epithelial cells, lung fibroblasts, and lung microvascular endothelial cells in 3D to generate airway organoids^28^. Under these conditions, randomly-seeded mixed cell populations underwent rapid condensation to self-organize into discrete epithelial and endothelial structures that were stable up to four weeks of culture^28^. Hild and Jaffe have described a protocol for the culture of *bronchospheres* from primary human airway basal cells. Mature *bronchospheres* are composed of functional multi-ciliated cells, mucin-producing goblet cells, and airway basal cells^29^. Finally, Snoeck and colleagues generated lung bud organoids from human pluripotent stem cells that recapitulate fetal lung development^30^. However, none of these approaches allows continued, long-term expansion of airway epithelium in vitro limiting their practical use. We therefore set out to establish long-term culture conditions of human airway epithelial organoids that contain all major cell populations and allow personalized human disease modelling.

## Results

### Generation and characterization of human airway organoids

We collected macroscopically inconspicuous lung tissue from non-small-cell lung cancer (NSCLC) patients undergoing medically indicated surgery and isolated epithelial cells through mechanical and enzymatic tissue disruption (see Methods). Following our experience with generating organoids from other adult human tissues^31-35^, we embedded isolated cells in basement membrane extract (BME) and activated/blocked signalling pathways important for airway epithelium (Supplementary Table 1). Under optimized conditions, 3D organoids formed within several days (94% success rate, n=18) and could be passaged by mechanical disruption at 1:2 to 1:4 ratios every other week for at least one year. Accordingly, they proliferated at comparable rates regardless of passage number (Extended Data Fig. 1a).

Since organoids were composed of a polarized, pseudostratified airway epithelium containing basal, secretory, brush, and multi-ciliated cells (Fig. 1a, b, Supplementary Video 1), we termed them airway organoids (AOs). Within the organoids, cells that stained positive for basal cell marker Keratin-14 (KRT14), club cell marker secretoglobin family 1A member 1 (SCGB1A1), cilia marker acetylated α-tubulin, or goblet cell marker mucin 5AC (MUC5AC) localized to their corresponding *in vivo* positions (Fig. 1c, Extended Data Fig. 1b). Secretory cells as well as cilia were functional as evidenced by immunofluorescence analysis capturing secretion of SCGB1A1-positive mucus into the organoid lumen (Fig. 1c) and time lapse microscopy showing cilia whirling around secreted mucus (Supplementary Video 2, 3).

**Figure 1.**
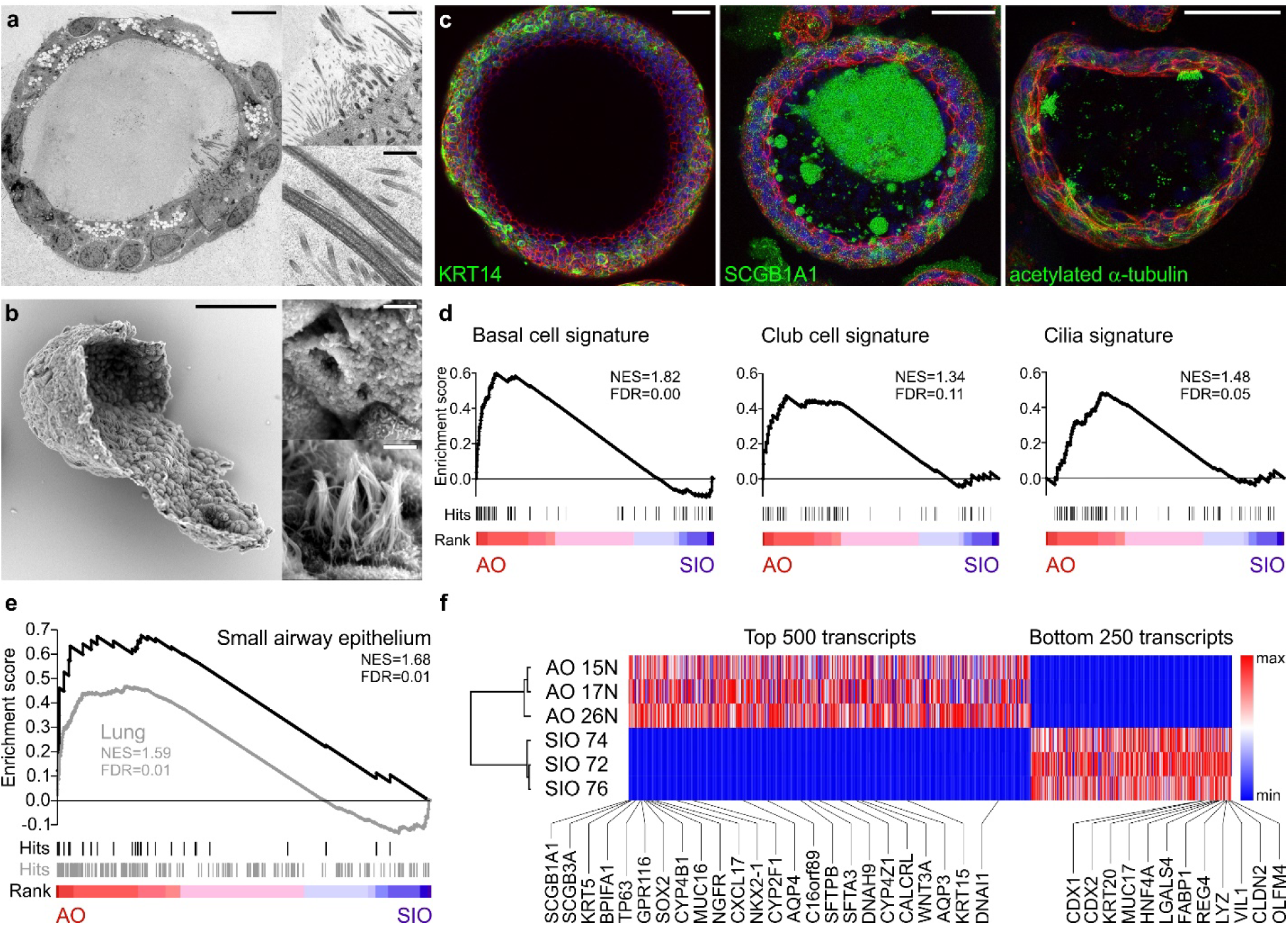
Characterization of airway organoids. **a**, Transmission electron micrograph of an AO cross-section showing the polarized, pseudostratified epithelium containing basal, secretory, brush, and multi-ciliated cells. Details display apical microvilli and cilia with their characteristic microtubule structure. Scale bars equal 10 μm, 2 μm, and 500 nm. See also Supplementary Videos 1-3. **b**, Scanning electron micrograph of a partially opened AO visualizing its 3D architecture, as well as basal and apical ultrastructure. Details display apical surfaces of secretory and multi-ciliated cells. Scale bars equal 50 μm (overview) and 2 μm (details). **c**, Immunofluorescent mid-sections of AOs showing markers for basal cells (KRT14), club cells (SCGB1A1), and cilia (acetylated α-tubulin). KRT14 is present exclusively in basally localized cells, SCGB1A1 stains luminal cells and multiple luminally secreted mucus clouds, and cilia cluster apically on luminal cells. Counterstained are the actin cytoskeleton (red) and nuclei (blue). Scale bars equal 50 μm. **d**, **e**, Gene set enrichment analysis plots showing strong enrichment of indicated gene signatures in transcriptomes of three AO lines compared to three small intestinal organoid (SIO) lines. NES = normalized enrichment score, FDR = false discovery rate. See Supplementary Table 2 for signatures and leading-edge genes. **f**, Hierarchical clustering of the indicated AO and SIO lines. Analysis is based on the top 500 and bottom 250 expressed genes in averaged AO vs SIO transcripts (full list in Supplementary Table 2). Gradients depict the relative maximum and minimum values per transcript. Names and ranks of signature lung and intestinal genes are indicated.

We next compared expression levels of selected genes in ten independently established AO lines vs whole lung tissue by quantitative PCR (qPCR) (Extended Data Fig. 1c). While equally expressing the general lung marker NKX2-1 and several airway specific markers, AOs expressed virtually no HOXA5 (a bona fide lung mesenchyme gene) or alveolar transcripts emphasizing their airway epithelial composition. To identify global AO gene expression and verify maintenance of airway identity upon extended culturing, we sequenced RNA of three high-passage AO lines and three small intestinal organoid (SIO) lines. Upon comparative transcriptome ranking, we performed gene set enrichment analyses (GSEA) using gene lists for specific lung cell types (see Supplementary Table 2 and Methods)^36-39^. Basal cell, club cell, and cilia signatures were strongly enriched in AOs (Fig. 1d), whereas alveolar and secretory cell signatures were not, the latter due to the presence of secretory cells in both AOs and SIOs (Extended Data Fig. 1c, d). Bulk lung as well as small airway epithelial signatures strongly correlated with AOs (Fig. 1e). Hallmark lung genes encoding for secretoglobins, cytochromes, p63, SOX2, and others were consistently among the highest AO enriched genes (Fig. 1f). High levels of WNT3A transcripts explained why AOs – in contrast to SIOs – did not require the addition of exogenous WNT3A to the culture media. Maximum WNT pathway activation caused upregulation of the proposed lung stem cell gene LGR6, while WNT pathway inhibition caused loss of LGR6 as well as of the generic epithelial stem cell marker LGR5 (Extended Data Fig.1e)^40,41^. Taken together, our culture conditions allow long term expansion of AOs that retain major characteristics of the *in vivo* epithelium.

### Airway organoids from patients with cystic fibrosis recapitulate central disease features and swell upon modulation of CFTR as well as activation of TMEM16A

Rectal organoids are being successfully used as functional model for cystic fibrosis (CF)^42^, a multi-organ disease with extensive phenotypic variability caused by mutations in the CF transmembrane conductance regulator gene (CFTR)^43^. Following opening of the CFTR channel by cAMP-inducing agents (e.g. forskolin), anions and fluid are transported to the organoid lumen resulting in rapid organoid swelling^44^, allowing personalized *in vitro* drug screenings^45^. Modelling of the primarily affected CF lung epithelium has relied on air-liquid interface cultures and is complicated by limited cell expansion and lengthy differentiation protocols^46^. To assess the AO approach for CF disease modeling, we applied forskolin and observed a dose-dependent swelling response that was largely, but not entirely, abrogated upon chemical inhibition of CFTR (Fig. 2a, Extended Data Fig. 2a), indicating the presence of additional ion channels. Indeed, AOs – but not rectal organoids – swell upon addition of E_act_ (Fig. 2b, Extended Data Fig. 2a) an activator of the chloride channel TMEM16A^47,48^. We next established five AO lines from fresh or cryopreserved broncho-alveolar lavage fluid of independent CF patients (62% success rate, n=8). CF AOs presented a much thicker layer of apical mucus compared to patient matched rectal organoids and wild-type AOs (Fig. 2c, Extended Data Fig. 2b), recapitulating the *in vivo* CF phenotype. Forskolin-induced swelling was dramatically reduced in CF compared to wild-type AOs, correlated with severity of tested CFTR genotypes^49^, and could be augmented with the CFTR modulators VX-770 and VX-809 (Fig. 2d, Extended Data Fig. 2c) in agreement with clinical data^30^. E_act_-stimulation caused a swelling response similar to stimulation with forskolin ± VX-770 and VX-809 (Fig. 2d, Extended Data Fig. 2c). Taken together, these experiments establish that AO lines derived from small amounts of patient material can recapitulate central features of CF, a classic example of monogenic diseases.

**Figure 2.**
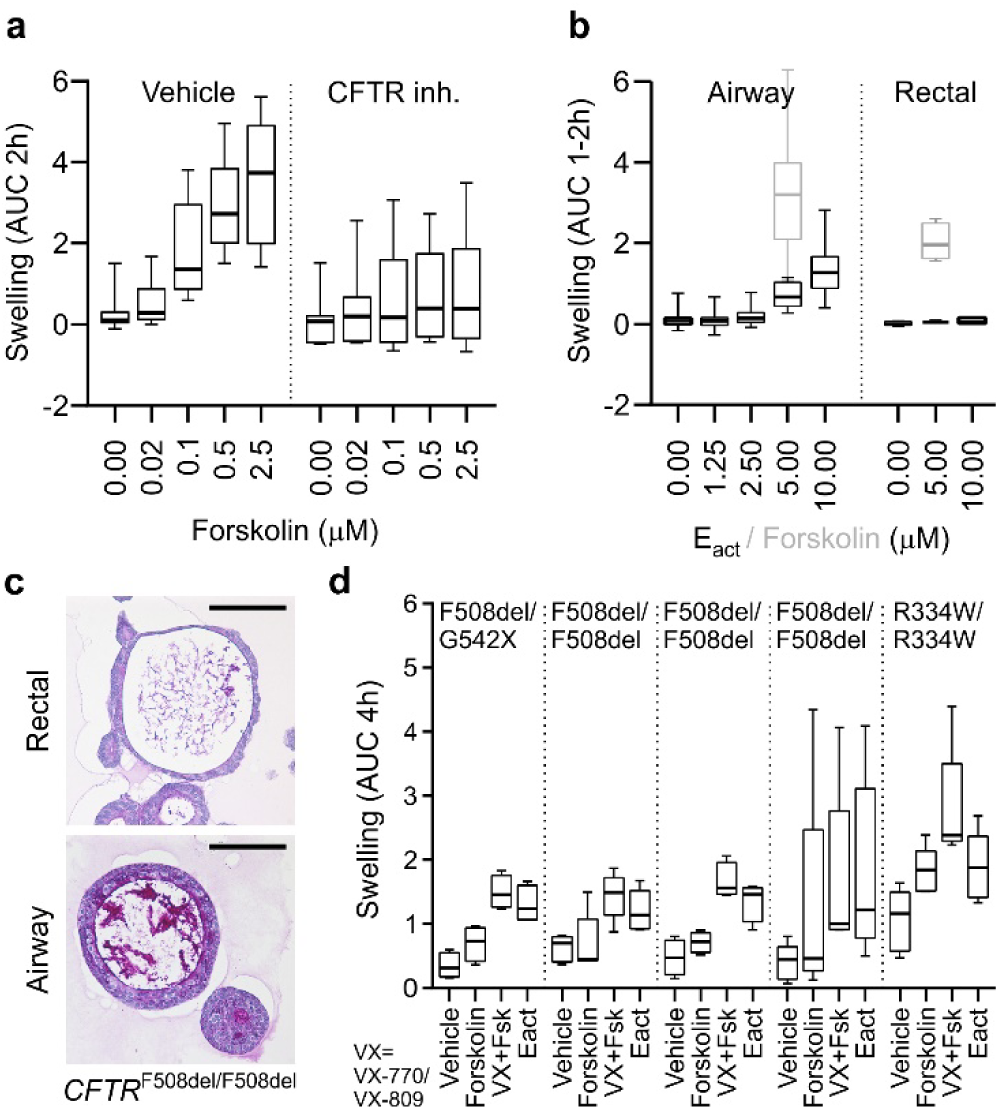
Airway organoids to study cystic fibrosis. **a**, Box-and-Whisker plot showing concentration-dependent forskolin-induced swelling of AOs in the absence and presence of CFTR inhibitors CFTRinh-172 and GlyH101. Upon CFTR inhibition, swelling is noticeably decreased but not absent. Shown are pooled data from three different AO lines used in each of three independent experiments. AUC = area under the curve. **b**, Box-and-Whisker plot showing concentration-dependent E_act_-induced swelling of AOs, but not rectal organoids (black outlines). Forskolin causes swelling in both organoid types (grey outlines). Shown are pooled data from three different AO and two different rectal organoid lines used in three to four independent experiments. Swelling was linear for 2h for AOs, but only 1h for rectal organoids. See Extended Data Fig. 2a for respective time course plots. **c**, Representative histological sections of periodic acid–Schiff (PAS) stained organoids from a CF patient with *CFTR*^F508del/F508del^ mutation. Note the thick layer of PAS-positive polysaccharides apically lining the airway epithelium. Rectal organoids were generated from rectal biopsies, AOs were generated from broncho-alveolar lavages (BALs). Scale bars equal 50 μm. See Extended Data Fig. 2b for PAS-stained wild-type and *CFTR*^R334W/R334W^ organoid sections. **d**, Box-and-Whisker plot showing swelling assays of several CF patient AO lines carrying the indicated CFTR mutations. Forskolin-induced swelling rarely exceeds vehicle controls in CF AOs, but increases in the presence of the CFTR-mutant modulating drugs VX-770 and VX-809. E_act_-induced swelling exceeds forskolin-induced swelling to a similar extend as pre-treatment with VX-770 and VX-809 in four out of five CF AO lines. Shown are pooled data of four to five independent experiments. See Extended Data Fig. 2c for selected time course plots.

### Generation of TP53 mutant airway organoids and selective expansion of tumoroids from primary and metastatic lung cancer

We next tested if AOs can be used to model lung cancer, a global respiratory disease burden^31,32^. We have previously observed that remnants of normal epithelium in carcinoma samples will rapidly overgrow tumor tissue^33,35^. While we successfully generated organoid lines in the majority of cases (88% success rate, n=16), we could not selectively expand lung tumoroids by removing a single medium component (such as Wnt3A in colon tumoroids^35^), due to the diversity of mutated signalling pathways in lung cancer^51^. We reasoned Nutlin-3a^53^ could drive *TP53* wild-type AOs into senescence or apoptosis and allow outgrowth of tumoroids with mutant p53, present in a large proportion of NSCLCs^54^. As proof of concept, we generated frameshift mutations in *TP53* of wild-type AOs with CRISPR-Cas9, causing Nutlin-3a resistance in selected sub-clones (Fig. 3a, Extended Data Fig. 3) due to lost p53 protein (Fig. 3b). Using Nutlin-3a selection, we indeed generated pure lung tumoroid lines from several NSCLC subtypes (including adenocarcinoma and large cell carcinoma) recapitulating fundamental histological characteristics of the respective primary tumors (Fig. 3c). In addition, we established pure tumoroid lines from needle biopsies of metastatic NSCLCs circumventing the need for Nutlin-3 selection (Fig. 3c, d) due to the absence of normal lung tissue (28% success rate, n=18). Hotspot DNA sequencing of selected cancer genes in tumoroids revealed loss- (e.g. *STK11*, *TP53*) and gain-of-function mutations (e.g. *KRAS*, *ERBB2*) (Fig. 3d, Supplementary Table 3). In conclusion, we show that AOs are amenable to subcloning and gene editing and we provide basic protocols for the selective outgrowth of p53 mutant TAOs from primary and metastatic NSCLCs.

**Figure 3.**
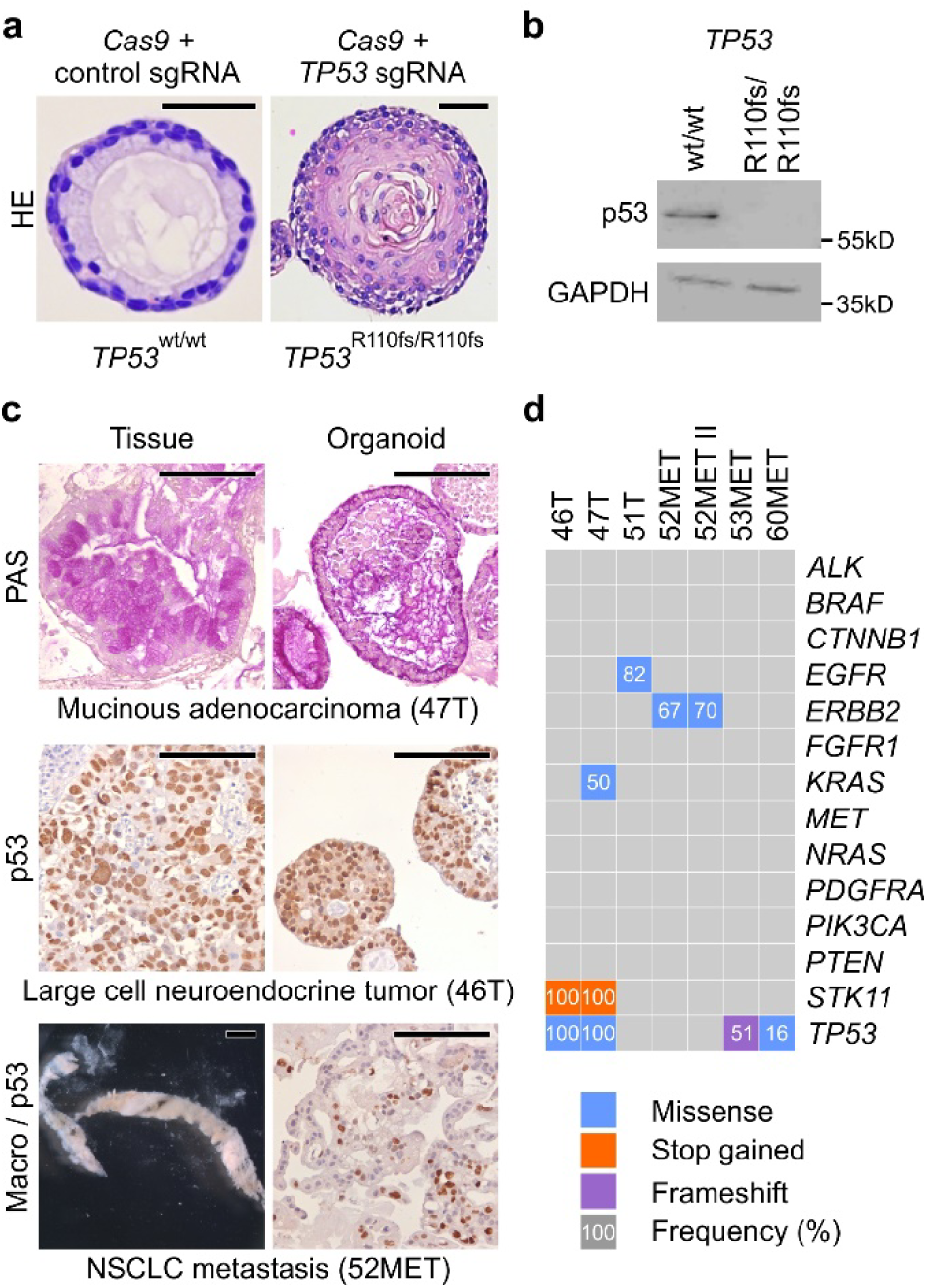
Modelling lung cancer using airway organoids. **a**, Histology of representative control and engineered *TP53*^R110fs/R110fs^ AO clones. Scale bars equal 100 μm. **b**, Western blot analysis of *TP53*^wt^ and *TP53*^R110fs/R110fs^ AOs show absence of p53 in the latter. **c**, Examples of TAOs derived from different resected primary lung cancer types (top and middle) as well as from a biopsy of metastatic NSCLC (bottom). Note the preservation of histological features between tissue-organoid pairs including PAS-positive mucus deposits, nuclear and cellular size abnormalities, and p53 immunolabelling. Scale bars equal 200 μm (histology) and 2 mm (macro). **d**, Mutation status of selected lung cancer genes in several TAO lines derived from primary as well as metastatic lung cancer. See Supplementary Table 3 for details. Note the preservation of mutant ERBB2 frequencies in 52MET and 52MET II TAOs, which have been derived from independent biopsies of the same cancer three months apart.

### RSV infection causes dramatic epithelial remodelling in airway organoids

Respiratory infections pose an even bigger global disease burden than genetic lung diseases^52^. RSV infections alone cause millions of annual hospital admissions and hundreds of thousands of deaths among young children^55^ due to bronchiolitis, oedema, and abnormalities of the airway epithelium (necrosis, sloughing)^56^. Since disease pathology is incompletely understood, we tested if AOs can serve as *in vitro* model for RSV infection. RSV replicated readily in multiple AO lines (Fig. 4a, Supplementary Video 4) but failed to do so after pre-incubation with commercial palivizumab (an antibody against the RSV F-glycoprotein that prevents RSV-cell fusion) indicating specific virus-host interaction (Fig. 4b)^57^. Morphological analysis of RSV-infected AOs revealed massive epithelial abnormalities (Fig. 4c) that recapitulated *in vivo* phenomena including cytoskeletal rearrangements, apical extrusion of infected cells, and syncytia formation (Fig. 4c, d). We used time-lapse microscopy to visualize the underlying dynamics and surprisingly found RSV-infected AOs to rotate and move through BME to fuse with neighbouring AOs (Fig. 4e, Supplementary Video 4). Organoid motility was caused by increased motilities of all cells within RSV-infected AOs (Fig. 4f, Supplementary Videos 5-6). Mathematical modelling and simulation suggested organoid rotation to be the macroscopic manifestation of coordinated cell motilities (Extended Data Fig. 4, Supplementary Video 7). RNA-sequencing of RSV-vs mock-infected AOs showed strong enrichment for genes involved in interferon α/β signalling (particularly cytokines), which correlated with enrichment of migratory as well as of antiviral response genes (Fig. 4g, h, Supplementary Table 4). Enzyme-linked immunosorbent assays confirmed the secretion of significant quantities of cytokines such as IP-10 and RANTES from RSV-infected AOs (Fig. 4i). To test whether secreted cytokines attract neutrophils (known to accumulate at RSV-infection sites *in vivo^58^)*, we co-cultured RSV-infected AOs with freshly isolated primary human neutrophils. Time-lapse microscopy indeed showed preferential neutrophil recruitment to RSV-infected AOs compared to mock controls (Fig. 4j, Supplementary Videos 8 and 9) allowing experimental dissection of immune cell interaction with RSV-infected airway epithelium *ex vivo*. Mechanistically, the non-structural protein NS2 has been shown to induce the shedding of dead cells *in vivo*, leading to airway obstructions as well as to be required for efficient propagation of the virus *in vivo*^59,60^. RSV^ΔNS2^ indeed replicated less efficiently than RSV^wt^ in AOs (Fig. 4k, Supplementary Video 10), while inducible overexpression of NS2 alone in non-infected AOs induced motility and fusion of RSV-infected AOs, underlining the crucial role of the protein in the modification of the behaviour of the infected cell (Fig. 4l, Extended Data Fig. 5, Supplementary Video 11). Taken together, RSV-infected AOs recapitulate central disease features and reveal their underlying highly dynamic epithelial remodelling.

**Figure 4.**
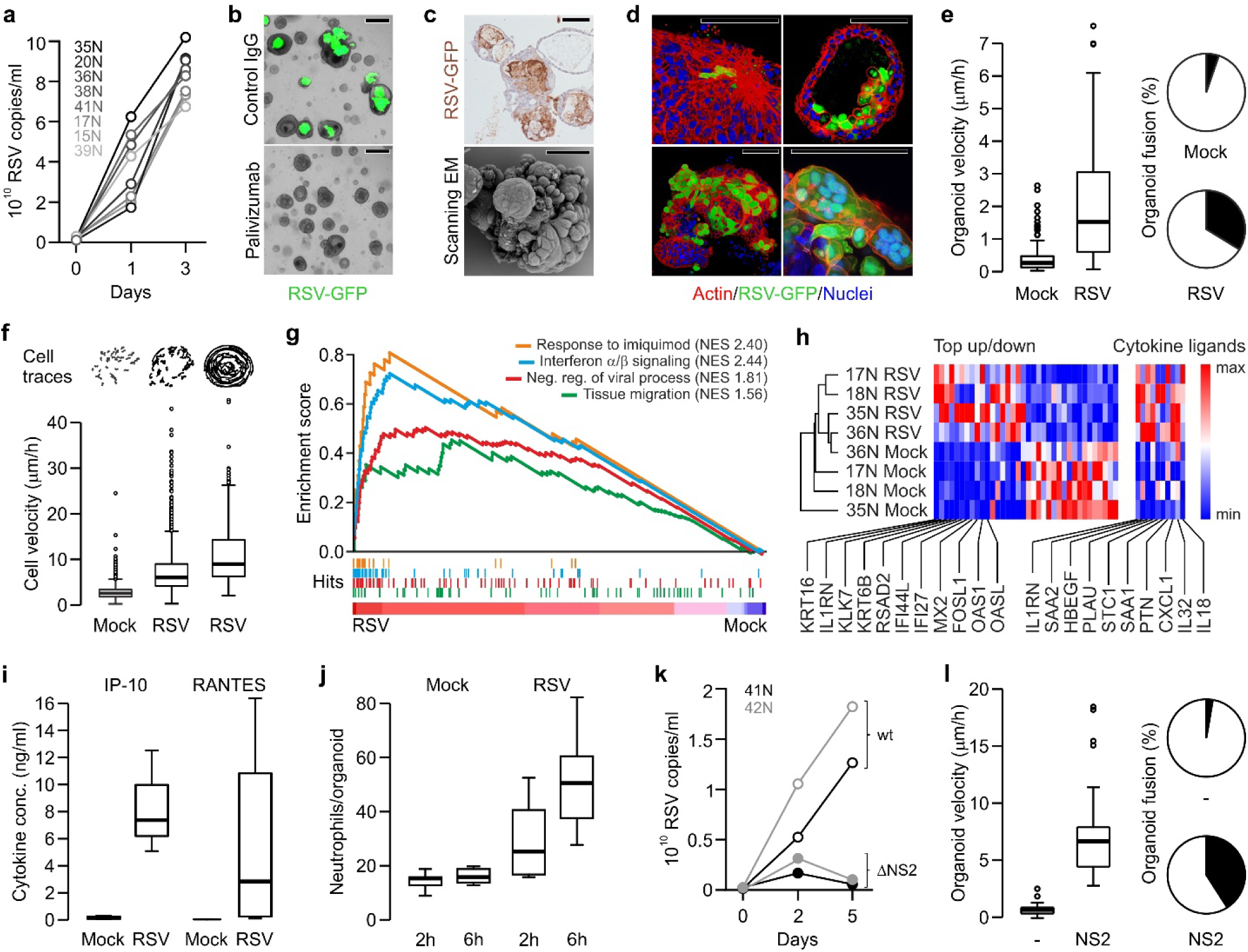
Modelling RSV infection with airway organoids. **a**, Quantitative PCR showing RSV replication kinetics in the indicated AO lines. **b**, Representative phase contrast/GFP overlays of RSV-infected AOs (5d post infection). GFP signal is absent following pre-incubation with palivizumab (fusion blocking antibody) but not control IgG. Scale bars equal 100 μm. **c**, Immunohistochemical RSV-staining (top) and scanning electron micrograph (bottom) of RSV-infected AOs (5d post infection) show organoid fusion and blebbing. Scale bars equal 50 μm. **d**, 3D reconstructions of immunolabeled AOs 3d after RSV-infection showing (clockwise) cytoskeletal rearrangements, apical extrusion of infected cells, syncytia formation, and organoid fusion. Scale bars equal 50 μm. **e**, RSV infection causes increased organoid velocity (Box-and-Whisker plot) and organoid fusion (pie chart). See also Supplementary Video 4. **f**, Cells within RSV-infected organoids are more motile than cells in control organoids irrespective of induced organoid rotation. Cell traces above individual Box-and-Whisker plots are from representative organoids. See also Extended Data Fig. 4 and Supplementary Videos 5-6. **g**, GSEA plots showing strong enrichment of indicated gene signatures in transcriptomes of four independently RSV- vs Mock-infected AO lines. NES = normalized enrichment score. See Supplementary Table 4 for signatures and leading-edge genes. **f**, Hierarchical clustering of the indicated AO lines displaying the 36 most differentially expressed genes of RSV- vs Mock-infected AO transcripts as well as selected cytokines (see Supplementary Table 4). Gradients depict the relative maximum and minimum values per transcript. Names and ranks of selected genes involved in migration (KRT16, KRT6B), interferon signaling (IL1RN, IFI44L), and viral response (MX2, OAS1) are indicated. **i**, ELISA-based quantification of cytokines secreted by Mock- vs RSV-infected AOs. **j**, Box-and-Whisker plot showing increased numbers of primary human neutrophils populating RSV-compared to Mock-infected AOs. See also Supplementary Videos 8 and 9. **k**, Quantitative PCR showing replication kinetics of wild-type (wt) and mutant RSV lacking NS2 (ΔNS2) in the indicated AO lines. See also Supplementary Video 10. **l**, Inducible overexpression of NS2 causes increased organoid velocity (Box-and-Whisker plot) and fusion (pie chart). See Extended Data Fig. 5, Supplementary Video 11.

## Discussion

Here we describe a versatile approach to establishing adult human airway epithelial organoids, containing all major cellular elements. We show the relative ease with which these can be grown from small amounts of routinely obtained patient material and provide evidence that the technology allows long-term expansion of such organoids from healthy individuals, but also from patients with a hereditary (CF) or malignant lung disease (NSCLC). We exploit the potential to derive sub-clones from AOs to demonstrate the feasibility of CRISPR gene editing.

Finally, we show that AOs readily allow modelling of viral infections such as RSV and for the first time demonstrate the possibility to study neutrophil-epithelium interaction in an organoid model. Taken together, we anticipate that human AOs will find broad applications in the study of adult human airway epithelium in health and disease.

## Methods

### Procurement of human material and informed consent

The collection of patient data and tissue for the generation and distribution of airway organoids has been performed according to the guidelines of the European Network of Research Ethics Committees (EUREC) following European, national, and local law ^1^. In the Netherlands, the responsible accredited ethical committees reviewed and approved the studies in accordance with the *‘Wet medisch-wetenschappelijk onderzoek met mensen’* (medical research involving human subjects act)^2^. The medical ethical committee UMC Utrecht (METC UMCU) approved protocols 07-125/C (isolation and research use of neutrophils from healthy donors), TCBio 15-159 (isolation and research use of broncho-alveolar lavage fluid of CF patients), and TCBio 14-008 (generation of organoids from rectal biopsies of CF patients). The *‘Verenigde Commissies Mensgebonden Onderzoek’* of the St. Antonius Hospital Nieuwegein approved protocol Z-12.55 (collection of blood, generation of normal and tumor organoids from resected surplus lung tissue of NSCLC patients). The Medical Ethics Committees of the Netherlands Cancer Institute Amsterdam approved PTC14.0929/M14HUP (collection of blood, generation of normal and tumor organoids from resected surplus lung tissue of NSCLC patients), and PTC14.0928/M14HUM (generation of tumor organoids from biopsies of metastatic NSCLC). All patients participating in this study signed informed consent forms approved by the responsible authority. In all cases, patients can withdraw their consent at any time, leading to the prompt disposal of their tissue and any derived material. AOs established under protocols Z-12.55, PTC14.0929/M14HUP, and PTC14.0928/M14HUM were biobanked through Hubrecht Organoid Technology (HUB, www.hub4organoids.nl). Future distribution of organoids to any third (academic or commercial) party will have to be authorized by the METC UMCU/TCBio at request of the HUB in order to ensure compliance with the Dutch medical research involving human subjects’ act.

### Tissue processing

Solid lung tissue was minced, washed with 10 ml AdDF+++ (Advanced DMEM/F12 containing 1x Glutamax, 10 mM HEPES, and antibiotics) and digested in 10 ml airway organoid medium (Supplementary Table 1) containing 1-2 mg⋅ml^-1^ collagenase (Sigma-C9407) on an orbital shaker at 37°C for 1-2 h. The digested tissue suspension was sequentially sheared using 10 ml and 5 ml plastic and flamed glass Pasteur pipettes. After every shearing step the suspension was strained over a 100 μm filter with retained tissue pieces entering a subsequent shearing step with ~10 ml AdDF+++. 2% FCS was added to the strained suspension before centrifugation at 400 rcf. The pellet was resuspended in 10ml AdDF+++ and centrifuged again at 400 rcf. In case of a visible red pellet, erythrocytes were lysed in 2 ml red blood cell lysis buffer (Roche-11814389001) for 5 min at room temperature before the addition of 10 ml AdDF+++ and centrifugation at 400 rcf. Broncho-alveolar lavage fluid was collected and incubated with 10 ml airway organoid medium containing 0.5 mg⋅ml^-1^ collagenase (Sigma-C9407), 0.5% (w⋅v^-1^) Sputolysin (Boehringer Ingelheim-SPUT0001), 250 ng⋅ml^-1^ amphotericin B, and 10 μg⋅ml^-1^ gentamicin on an orbital shaker at 37°C for 10-30 min. Next, the suspension was mildly sheared using a 1 ml pipet tip and strained over a 100 μm filter. 2% FCS was added to the strained suspension before centrifugation at 400 rcf. The pellet was resuspended in 10ml AdDF+++ and centrifuged again at 400 rcf.

### Organoid culture

Lung cell pellets were resuspended in 10 mg⋅ml^-1^ cold Cultrex growth factor reduced BME type 2 (Trevigen-3533-010-02) and 40 μl drops of BME-cell suspension were allowed to solidify on prewarmed 24-well suspension culture plates (Greiner-M9312) at 37°C for 10-20 min. Upon completed gelation, 400 μl of AO medium were added to each well and plates transferred to humidified 37°C / 5% CO_2_ incubators at ambient O_2_. Medium was changed every 4 days and organoids were passaged every 2 weeks: cystic organoids were resuspended in 2 ml cold AdDF+++ and mechanically sheared through flamed glass Pasteur pipettes. Dense (organoids were dissociated by resuspension in 2 ml TrypLE Express (Invitrogen-12605036), incubation for 1-5 min at room temperature, and mechanical shearing through flamed glass Pasteur pipettes. Following the addition of 10 ml AdDF+++ and centrifugation at 300 rcf or 400 rcf respectively, organoid fragments were resuspended in cold BME and reseeded as above at ratios (1:1 – 1:6) allowing the formation of new organoids. Single cell suspensions were initially seeded at high density and reseeded at a lower density after ~1 week. NSCLC organoids could be distinguished from normal regular cystic organoids by morphology (size, irregular shape, thick organoid walls, dense) as well as histology. Separation from normal AOs was achieved by manual separation and in case of *TP53* mutations by the addition of 5 μM Nutlin-3 (Cayman Chemicals-10004372) to the culture medium. Intestinal organoids were cultured as previously described ^3^.

### Luminescent viability assay

Single cell suspensions from AOs were generated as described and counted. 3⋅10^3^ cells per replicate were plated in BME and incubated in AO medium as described. At the indicated time points, the medium was removed and organoids were lysed in CellTiter Glo 3D (Promega) according to the manufacturer’s instructions. Luminescence was measured using a microplate luminometer (Berthold Technologies).

### Electron microscopy

For transmission EM, organoids were placed in BME on 3 mm diameter and 200 μm depth standard flat carriers for high pressure freezing and immediately cryoimmobilized using a Leica EM high-pressure freezer (equivalent to the HPM10), and stored in liquid nitrogen until further use. They were freeze-substituted in anhydrous acetone containing 2% osmium tetroxide and 0.1% uranyl acetate at –90°C for 72 hours and warmed to room temperature at 5°C per hour (EM AFS-2, Leica). The samples were kept for 2 h at 4°C and 2 h at room temperature. After several acetone rinses (4x 15 min), samples were infiltrated with Epon resin during 2 days (acetone: resin 3:1 3 h; 2:2 – 3 h; 1:3 – overnight; pure resin-6 h + overnight + 6 h + overnight + 3 h). Resin was polymerised at 60°C during 48 hours. Ultrathin sections from the resin blocks were obtained using an UC6 ultramicrotome (Leica) and mounted on Formvar-coated copper grids. Grids were stained with 2% uranyl acetate in water and lead citrate. Sections were observed in a Tecnai T12 Spirit electron microscope equipped with an Eagle 4kx4k camera (FEI Company) and large EM overviews were collected using the principles and software described earlier ^4^. For scanning EM, organoids were removed from BME, washed with excess AdDF+++, fixed for 15 min with 1% (v/v) glutaraldehyde (Sigma) in phosphate buffered saline (PBS) at room temperature, and transferred onto 12 mm poly-L-lysine coated coverslips (Corning). Samples were subsequently serially dehydrated by consecutive 10 min incubations in 2 ml of 10% (v/v), 25% (v/v) and 50% (v/v) ethanol-PBS, 75% (v/v) and 90% (v/v) ethanol-H_2_O, and 100% ethanol (2x), followed by 50% (v/v) ethanol-hexamethyldisilazane (HMDS) and 100% HMDS (Sigma). Coverslips were removed from the 100% HMDS and air-dried overnight at room temperature. Organoids were manipulated with 0.5 mm tungsten needles using an Olympus SZX9 light microscope and mounted onto 12 mm specimen stubs (Agar Scientific). Following gold-coating to 1 nm using a Q150R sputter coater (Quorum Technologies) at 20 mA, samples were examined with a Phenom PRO table-top scanning electron microscope (Phenom-World).

### Time-lapse microscopy

Bright-field AO time-lapse movies were recorded at 37°C and 5% CO_2_ on an AF7000 microscope equipped with a DFC420C camera using LAS AF software (all Leica). Bright-field cilia movement was recorded using the same set-up equipped with a Hamamatsu C9300-221 high speed CCD camera (Hamamatsu Photonics) at 150 frames per second using Hokawo 2.1 imaging software (Hamamatsu Photonics). Confocal imaging was performed using the following microscopes at 37°C and 5% CO_2_: SP8X (Leica), LSM710, LSM800 (both Zeiss), and Ultraview VoX spinning disk (Perkin Elmer).

### Fixed immunofluorescence microscopy and immunohistochemistry

Organoids were removed from BME, washed with excess AdDF+++, fixed for 15 min in 4% paraformaldehyde, permeabilized for 20 min in 0.2% Triton X-100 (Sigma), and blocked for 45 min in 1% BSA. Organoids were incubated with primary antibodies over night at 4°C (anti-Keratin 14, Biolegend 905301; anti-SCGB1A1, Santa Cruz sc-9773; anti-acetylated α-tubulin Santa Cruz sc-23950; anti-Mucin 5AC, Santa Cruz sc-21701), washed three times with PBS, incubated with secondary antibodies (Invitrogen) over night at 4°C, washed two times with PBS, incubated with indicated additional stains (DAPI, life technologies D1306, phalloidin-atto 647, Sigma 65906), washed two times with PBS, and mounted in VECTASHIELD hard-set antifade mounting medium (Vectorlabs). Samples were imaged on SP5 and SP8X confocal microscopes using LAS X software (all Leica) and processed using ImageJ. For histological analysis, tissue and organoids were fixed in 4% paraformaldehyde followed by dehydration, paraffin embedding, sectioning, and standard HE and PAS stainings. Immunohistochemistry was performed using antibody against P53 (Santa Cruz, sc-126), GFP (Life technologies, A11122), and RSV (Abcam, ab35958). Images were acquired on a Leica Eclipse E600 microscope and processed using the Adobe Creative Cloud software package.

### RNA sequencing

AOs derived from three independent human donors were collected 6 days after splitting (at passage 16, 18 and 19, respectively). Small intestinal organoids were also derived from three different human donors and collected 2 days after splitting (passage 7). Total RNA was isolated from the collected organoids using RNeasy kit (QIAGEN) according to manufacturer’s protocol including DNaseI treatment. Quality and quantity of isolated RNA was checked and measured with Bioanalyzer2100 RNA Nano 6000 chips (Agilent, Cat. 5067-1511). Library preparation was started with 500ng of total RNA using the Truseq Stranded Total RNA kit with Ribo-Zero Human/Mouse/Rat set A and B by Illumina (Cat. RS-122-2201 and RS-122-2202). After preparation, libraries were checked with Bioanalyzer2100 DNA High Sensitivity chips (Cat. 5067-4626) as well as Qubit (Qubit^®^ dsDNA HS Assay Kit, Cat. Q32854), all samples had a RIN value of 10. Libraries were equimolarly pooled to 2 nM and sequenced on the Illumina Nextseq, 2×75bp high output (loaded 1.0-1.4 pM of library pools). Samples were sequenced to an average depth of 9.4 million mapped reads (SD 2.6 million). After sequencing, quality control, mapping and counting analyses were performed using our in-house RNA analysis pipeline v2.1.0 (https://github.com/CuppenResearch/RNASeq), based on best practices guidelines (https://software.broadinstitute.org/gatk/guide/article?id=3891). In short, sequence reads were checked for quality by FastQC (v0.11.4) after which reads were aligned to GRCh37 using STAR (v2.4.2a) and add read groups using Picard perform quality control on generated BAM files using Picard (v1.141). Samples passing QC were then processed to count reads in features using HTSeq-count (v0.6.1). Read counting for genes (ENSEMBL definitions GRCh37, release 74) resulted in a 12 sample count matrix with Ensembl gene identifiers. DeSeq (v1.18.0) was used for read normalisation and differential expression analysis. Gene set enrichment analysis (GSEA) was performed using signature gene lists for airway basal cells ^5^, club cells ^6^, cilia ^5^, small airway epithelial cells ^7^, and lung ^8^ against normalized RNA-seq reads of three AO and three SIO lines using GSEA software v3.0 beta2 ^9,10^. Normalized RNA-seq reads were averaged per organoid type and sorted by the log_2_ transformed ratio of AO over SIO. The 500 highest and 250 lowest transcripts were plotted using Morpheus (https://software.broadinstitute.org/morpheus) and CorelDraw X7.

### RNA preparation and qRT-PCR

Total RNA was isolated from three independent lung organoid stains after 6, 12, 24, and 48 hrs upon either Wnt activation or Wnt inhibition by using RNeasy kit (QIAGEN). To inhibit Wnt signalling, 6 days after splitting the culture media of lung organoids was changed to media lacking R-spondin and included IWP2 (3 μM, Stemgent). To activate Wnt signalling, Chir (3 μM) was added to the culture media 6 days after splitting. cDNA was synthesized from 1 μg of total RNA using GoScript (Promega). Quantitative PCR was performed in triplicate using the indicated primers, SYBR green, and BioRad systems. Gene expression was quantified using the ΔΔCt method and normalized by HPRT.

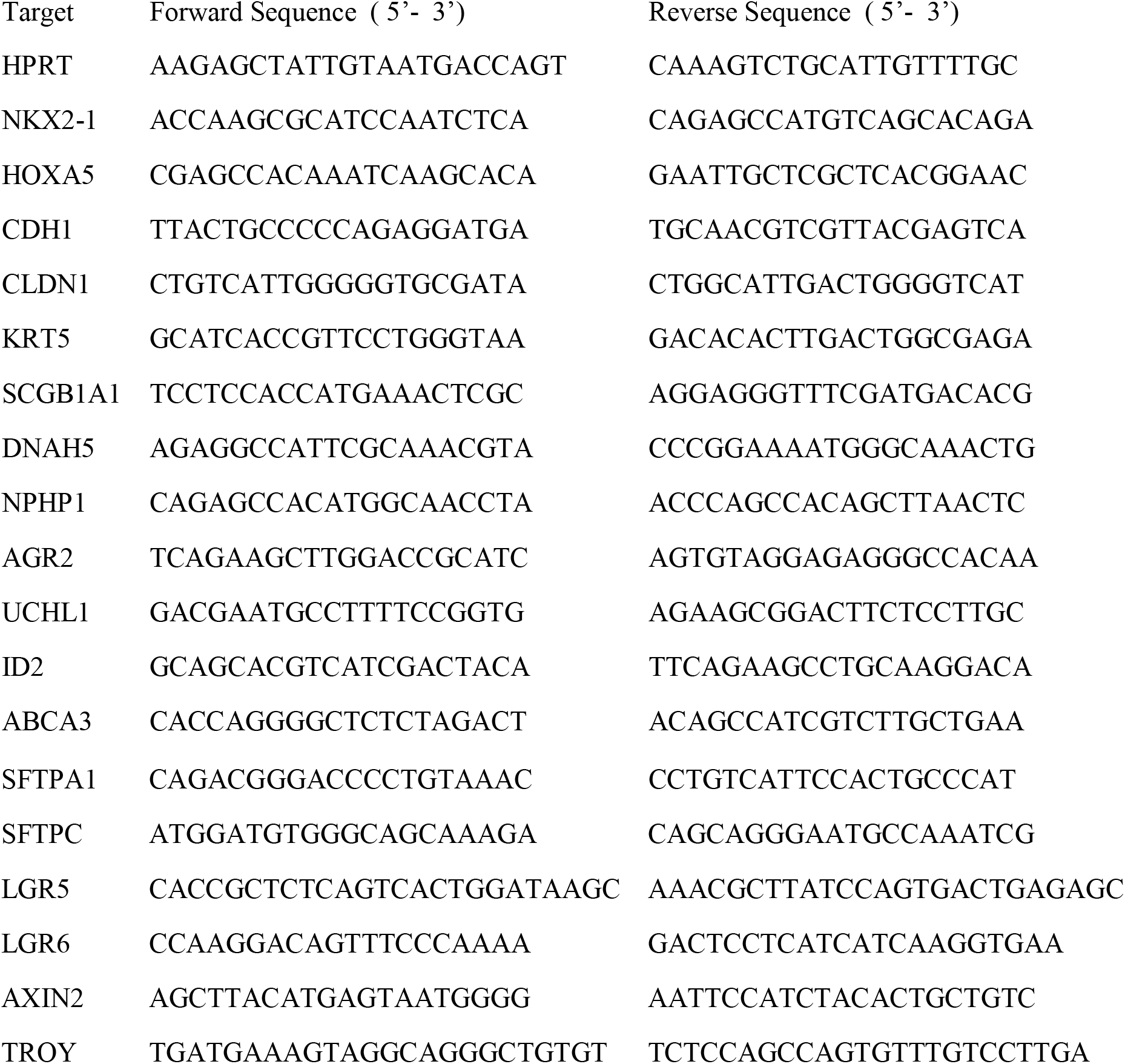

### Functional organoid swelling assay

Organoid swelling assays were performed as previously described ^11^. Intestinal organoids were cultured for 7 days, collected, and disrupted. Smallest fragments were selected and seeded for swelling assays. AOs were sheared, passed through a 70 μm strainer and cultured for 4 days. Organoids were harvested and seeded for swelling assays in 96-well plates (Greiner). We seeded 50-100 organoids in 5 μl drops (50% BME) per well overlaid with 100 μl culture medium after gel solidification. Cultures were incubated for 24 h with or without CFTR-targeting drugs VX-809 and VX-770 (3 μM and 1 μM, respectively, Selleck Chemicals). The next day, cultures were pre-incubated for 3 h with CFTR-inhibitors (150 μM CFTRinh-172 and 150 μM GlyH101, Cystic Fibrosis Foundation Therapeutics) as indicated. Calcein green (3 μM, Invitrogen) was added 1 h prior to stimulation with forskolin (Selleck Chemicals), E_act_ (Calbiochem), or DMSO (Sigma) at the indicated concentrations. Organoids were imaged per well (every 10 min, for 60- 120 min in total or every 30 min, for 270 min in total) using confocal live cell microscopy (Zeiss LSM710 and LSM800) at 37°C and 5% CO_2_. Data was analyzed using Volocity 6.1.1 (Perkin Elmer), Zen Blue (Zeiss) and Prism 5 and 6 (GraphPad).

### TP53 editing

The organoid lipofection protocol was followed as previously described ^12,13^ with a few adaptations. Briefly, AOs grown in full organoid media were trypsinized for 20 min at 37°C and sheared with a glass pipette to produce single cells. After dissociation, cells were resuspended in 450 μl growth medium and plated in 48-well plates at a confluency of >90%. For each transfection, a total of 1.5 μg of plasmid DNA (sgRNA, Cas9) in 50 μl of serum-free medium (Opti-MEM; Gibco) were mixed with 4 μl of Lipofectamine 2000 (Invitrogen) in 50 μl of serum-free medium making up a total volume of 100 μl, which was then added to the cells. The plate was centrifuged at 600g at 32^°^C for 1 h and incubated for 4-5 h at 37^°^C before the cells were embedded in Basement Membrane Extract (BME;Amsbio) in full organoid medium. 10 μM of Nutlin-1 (Cayman Chemicals) were added to the cultures 1 day after transfection and maintained for 16-20 days for mutant *P53* selection. For clonal expansion, single organoids were picked. At least two human AO lines were used for engineering mutant *P53* organoids. For sequencing, genomic DNA was isolated using Viagen Direct PCR (Viagen). Primers for PCR amplification using GoTaq Flexi DNA polymerase (Promega) were: *P53*_for, 5′- CAGGAAGCCAAAGGGTGAAGA-3′, *P53*_rev, 5′-CCCATCTACAGTCCCCCTTG-3′. PCR products were purified using the QIAquick PCR purification kit (Qiagen) and cloned into pGEM-T Easy vector system I (Promega), followed by sequencing using the T7 primer. The sgRNA *P53* sequence was specifically designed to target exon 3 of the *P53* gene and was: 5’- GGGCAGCTACGGTTTCCGTCGUUUUA-3’

### Western blotting

Samples were lysed using SDS lysis buffer (50 mM Tris HCl pH 6.8, 2% SDS) supplemented with Complete Protease Inhibitors (Roche). Protein concentration was determined using Nanodrop Lite (Thermo Scientific) and equal amounts of protein were loaded on SDS-PAGE gels before transferring on PVDF membranes (Millipore). Membranes were blocked in 5% milk (diluted in 0.01% PBS-Tween) and incubated overnight with antibodies against: P53 (Santa Cruz, sc-126) and GAPDH (Abcam, ab-9485).

### Hot-spot sequencing

AO gDNA was isolated using the DNeasy blood & tissue kit (Qiagen). A minimum of 100ng gDNA was submitted for sequencing cancer hotspots of Ion AmpliSeq^TM^ Cancer Hotspot Panel v2Plus at the NGS Core of the UMC Utrecht. The following exons were (partially) covered:

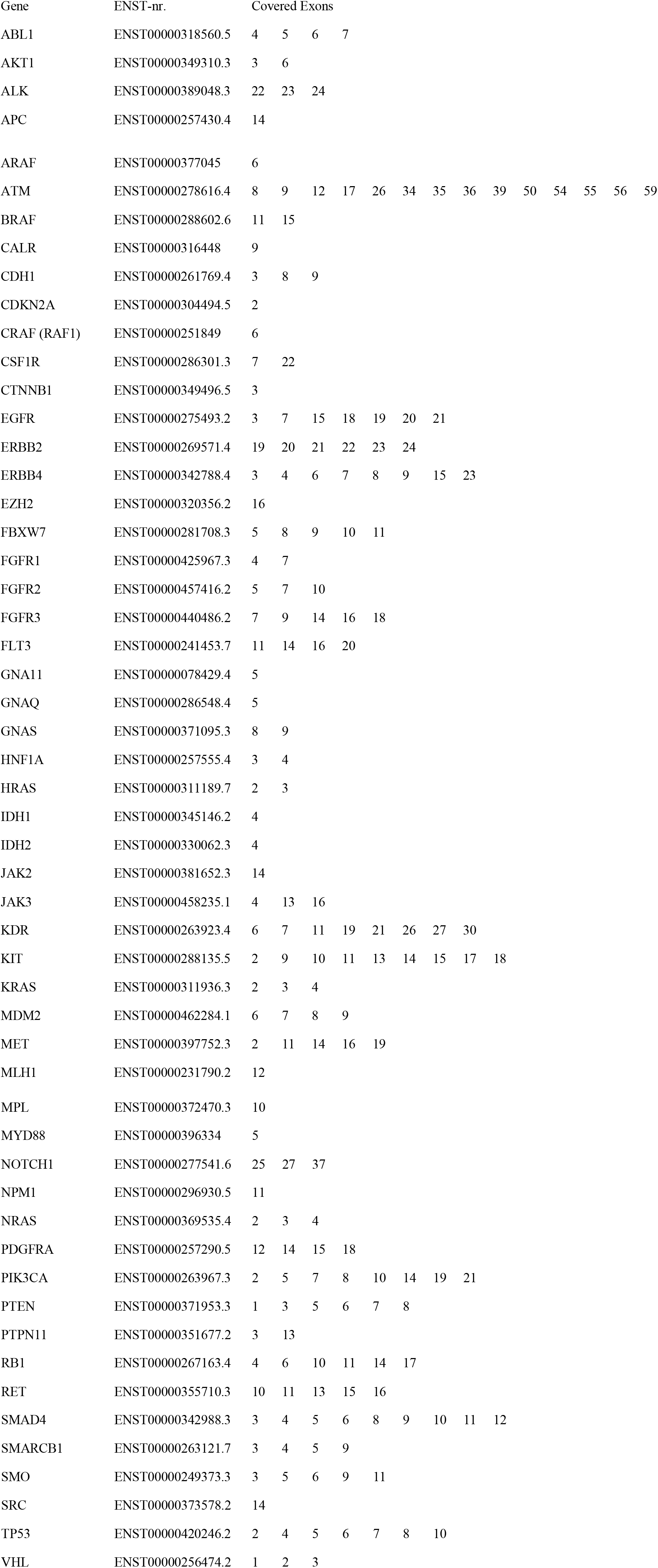

| Gene | ENST-nr. | Covered Exons |  |  |  |
| --- | --- | --- | --- | --- | --- |
| ABL1 | ENST00000318560.5 | 4 | 5 | 6 | 7 |
| AKT1 | ENST00000349310.3 | 3 | 6 |  |  |
| ALK | ENST00000389048.3 | 22 | 23 | 24 |  |
| APC | ENST00000257430.4 | 14 |  |  |  |

|  |  |  |  |  |  |  |  |  |  |  |  |  |  |  |  |  |  |
| --- | --- | --- | --- | --- | --- | --- | --- | --- | --- | --- | --- | --- | --- | --- | --- | --- | --- |
| ARAF | ENST00000377045 | 6 |  |  |  |  |  |  |  |  |  |  |  |  |  |  |  |
| ATM | ENST00000278616.4 | 8 | 9 | 12 | 17 | 26 | 34 | 35 | 36 | 39 | 50 | 54 | 55 | 56 | 59 | 61 | 63 |
| BRAF | ENST00000288602.6 | 11 | 15 |  |  |  |  |  |  |  |  |  |  |  |  |  |  |
| CALR | ENST00000316448 | 9 |  |  |  |  |  |  |  |  |  |  |  |  |  |  |  |
| CDH1 | ENST00000261769.4 | 3 | 8 | 9 |  |  |  |  |  |  |  |  |  |  |  |  |  |
| CDKN2A | ENST00000304494.5 | 2 |  |  |  |  |  |  |  |  |  |  |  |  |  |  |  |
| CRAF (RAF1) | ENST00000251849 | 6 |  |  |  |  |  |  |  |  |  |  |  |  |  |  |  |
| CSF1R | ENST00000286301.3 | 7 | 22 |  |  |  |  |  |  |  |  |  |  |  |  |  |  |
| CTNNB1 | ENST00000349496.5 | 3 |  |  |  |  |  |  |  |  |  |  |  |  |  |  |  |
| EGFR | ENST00000275493.2 | 3 | 7 | 15 | 18 | 19 | 20 | 21 |  |  |  |  |  |  |  |  |  |
| ERBB2 | ENST00000269571.4 | 19 | 20 | 21 | 22 | 23 | 24 |  |  |  |  |  |  |  |  |  |  |
| ERBB4 | ENST00000342788.4 | 3 | 4 | 6 | 7 | 8 | 9 | 15 | 23 |  |  |  |  |  |  |  |  |
| EZH2 | ENST00000320356.2 | 16 |  |  |  |  |  |  |  |  |  |  |  |  |  |  |  |
| FBXW7 | ENST00000281708.3 | 5 | 8 | 9 | 10 | 11 |  |  |  |  |  |  |  |  |  |  |  |
| FGFR1 | ENST00000425967.3 | 4 | 7 |  |  |  |  |  |  |  |  |  |  |  |  |  |  |
| FGFR2 | ENST00000457416.2 | 5 | 7 | 10 |  |  |  |  |  |  |  |  |  |  |  |  |  |
| FGFR3 | ENST00000440486.2 | 7 | 9 | 14 | 16 | 18 |  |  |  |  |  |  |  |  |  |  |  |
| FLT3 | ENST00000241453.7 | 11 | 14 | 16 | 20 |  |  |  |  |  |  |  |  |  |  |  |  |
| GNA11 | ENST00000078429.4 | 5 |  |  |  |  |  |  |  |  |  |  |  |  |  |  |  |
| GNAQ | ENST00000286548.4 | 5 |  |  |  |  |  |  |  |  |  |  |  |  |  |  |  |
| GNAS | ENST00000371095.3 | 8 | 9 |  |  |  |  |  |  |  |  |  |  |  |  |  |  |
| HNF1A | ENST00000257555.4 | 3 | 4 |  |  |  |  |  |  |  |  |  |  |  |  |  |  |
| HRAS | ENST00000311189.7 | 2 | 3 |  |  |  |  |  |  |  |  |  |  |  |  |  |  |
| IDH1 | ENST00000345146.2 | 4 |  |  |  |  |  |  |  |  |  |  |  |  |  |  |  |
| IDH2 | ENST00000330062.3 | 4 |  |  |  |  |  |  |  |  |  |  |  |  |  |  |  |
| JAK2 | ENST00000381652.3 | 14 |  |  |  |  |  |  |  |  |  |  |  |  |  |  |  |
| JAK3 | ENST00000458235.1 | 4 | 13 | 16 |  |  |  |  |  |  |  |  |  |  |  |  |  |
| KDR | ENST00000263923.4 | 6 | 7 | 11 | 19 | 21 | 26 | 27 | 30 |  |  |  |  |  |  |  |  |
| KIT | ENST00000288135.5 | 2 | 9 | 10 | 11 | 13 | 14 | 15 | 17 | 18 |  |  |  |  |  |  |  |
| KRAS | ENST00000311936.3 | 2 | 3 | 4 |  |  |  |  |  |  |  |  |  |  |  |  |  |
| MDM2 | ENST00000462284.1 | 6 | 7 | 8 | 9 |  |  |  |  |  |  |  |  |  |  |  |  |
| MET | ENST00000397752.3 | 2 | 11 | 14 | 16 | 19 |  |  |  |  |  |  |  |  |  |  |  |
| MLH1 | ENST00000231790.2 | 12 |  |  |  |  |  |  |  |  |  |  |  |  |  |  |  |

### RSV infection of AOs

rgRSV224 (GFP version), rrRSV (RFP version), rgRSV P-eGFP-M (RSV^wt^), and rgRSV∆NS2 13A ∆NS2 6120 P-eGFP-M (RSV^ΔNS2^) were produced as previously described ^14-16^. For infection, AOs were washed in cold AdDF+++ and sheared with a flamed Pasteur pipette. Per infection, 2 μl virus (~5⋅10^7^ pfu⋅ml^-1^) was placed in a U-bottom suspension 96-well and 50μl of sheared organoids (~500000 cells) in AdDF+++ were added. Empty wells were filled with PBS to prevent dehydration. Plates were incubated for 5 h at 37°C and 5% CO_2_. Afterwards, contents of the wells was taken up in AdDF+++ and washed three times with excess AdDF+++. Organoids were seeded as described before. Alternatively, individual AOs were microinjected with ~250 nl virus in AdDF+++ (~5⋅10^7^ pfu⋅ml^1^) using a micromanipulator and -injector (Narishige, M-152 and IM-5B) under a stereomicroscope (Leica, MZ75).

### Single-cell tracking

Cells were manually tracked by following the center of mass of their nuclei using custom written image analysis software. Random cells were selected on the initial time frame so that they uniformly covered the organoid surface. In infected organoids, particularly when rotating rapidly, a small fraction of cells could not be identified between some consecutive time frames due to their fast movement, limiting our ability to track the fastest-moving cells. In Fig. 4f and Extended Data Fig. 4b, three outliers were beyond the limits of the plot, reaching a maximum value of 117 μm⋅h-1. Due to the difficulty of tracking fast-moving cells, outliers above 50 μm⋅h-1 are likely underrepresented. For each cell at each time point, RFP intensity was calculated by summing the pixel intensities within a disk of a radius of 5 pixels, corresponding to 3.22 μm, on the XY plane around the center of mass of the nucleus. To distinguish between RFP^-^ and RFP^+^ cells, an intensity threshold was manually chosen so that all nuclei in non-infected organoids were categorized as RFP^-^.

### Mathematical modelling of collective cell motility

To understand how local interactions between individual cells could give rise to collective rotational movement on the level of the organoid, we built a mathematical model of cells migrating within the constraints of the tissue they are embedded in. In our model, cells move on the surface of a sphere to mimic the organoid geometry. Each cell has a polarity vector *p̄*_*I*_ that represents the direction of migratory force production. However, cells are constrained by adhesive connections to neighboring cells. In our model, these connections are modeled by harmonic springs. In general, due to these adhesive constraints the total force *f̄*_*i*_ acting on each cell has a different orientation than the intrinsic polarity *p̄*_*i*_. We assume that cells change their internal polarity over time to align with the direction of *f̄*_*i*_. Crucially, this interaction causes neighboring cells to align their movement. To reproduce the random motility observed in non-infected cells, we assume the polarity direction also fluctuates in a stochastic manner. This effect perturbs synchronization of movement between neighboring cells. Cells are initially distributed uniformly over the surface of sphere, with random orientation for their polarity vectors. The force on cell *i* is given by the sum of the propulsive force in the direction of the polarity *p̄*_*I*_ the spring forces *F̄*^*ij*^ due to the neighboring cells *j*:

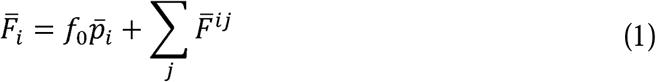

The spring force between a pair of cells at positions *r̄i*and *r̄j* is given by:

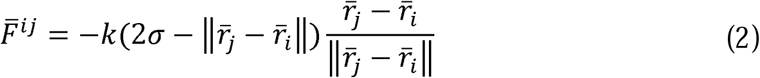
where *k* is the spring constant. Here, σ corresponds approximately to the radius of the cell and, hence, the spring force is attractive when it is extended beyond its natural length 2σ and repulsive when compressed. The model is then described fully by two sets of ordinary differential equations, for the cell positions *r*̄i and the cell polarities *p̄* _*i*_:

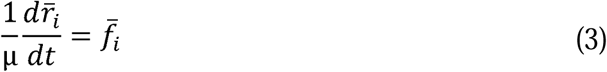

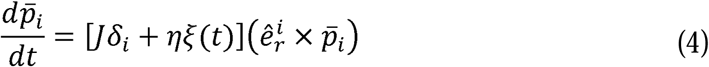

Here, cell movement is assumed to be overdamped, with μ the mobility. *f̄* _*i*_ is the projection of the total force *F̄* _*i*_ in the local plane of cell *i*, i.e. 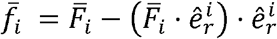 with 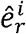 the unit vector orthogonal to the surface of the sphere at *r̄*_*i*_. The polarity vector *p̄*_*i*_ has length ||*p̄*_*i*_ ||=1and rotates with fixed angular velocity *J* in the direction that aligns it with *f̄*_*i*_. This direction is given by δ_*i*_ = cos *θ_i_*/|cos *θ_i_*| where θ_i_ is the angle between *p̄*_*i*_ and *f̄*_*i*_ and 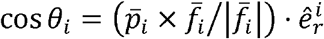 In addition, we incorporate intrinsic stochastic fluctuations to the direction of cell polarity by the noise term ξ(*t*), with noise strength *η*. In the absence of cell-cell communication, i.e. for *J*=0, the polarity vector *p̄*_*i*_ (*t*)will undergo a random walk that causes the polarity to deviate from its original polarity over time. Specifically, the correlation in polarity direction decreases in time as 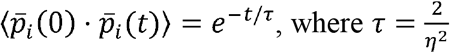 is the polarity persistence time.

### Simulation of the mathematical model

To generate random initial configurations of *N* particles distributed approximately equidistantly on a sphere with radius 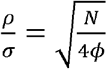, where, *ϕ* is the cellular packing fraction, particles were first placed at random positions on the sphere and then propagated using the spring force in Eq. (2) between nearest neighbors until global mechanical equilibrium was attained. At the start of the simulation, we assigned each particle *i* a polarity vector *p̄*_*i*_ with a random orientation in the tangent plane of the sphere at position *r*̄_*i*_ and determined the identity of the neighboring particles connected by a spring to particle *i* using Delauney triangulation. Equations (3) and (4) were integrated using the Euler-Maruyama method, with time-step *μk*⋅Δ*t*=5⋅10^−3^.To ensure that the position *r*̄_*i*_ and polarity vector *p̄*_*i*_ remain well-defined throughout the simulation, after each time step we projected *p̄*_*i*_ (*t*+Δ*t*)onto the tangent plane of the sphere at the new position *r*̄_*i*_(*t*+Δ*t*) and normalized the position and polarity so that||*r*̄_*i*_(*t*+Δ*t*)||=1and || *p̄*_*i*_(*t*+Δ*t*)||=1. Simulations were performed with *N* =100, and *ϕ*=1, and 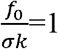 while we varied the parameters (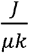) and (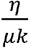) While we use dimensionless parameters here, in the main text we use, without any loss of generality, parameters values for *μ,k,*σ=1.

### Multiplex Immunoassays

AOs were infected with/without RSV as described. 3d after infection equal numbers of AOs were released from Matrigel using Cell Recovery Solution (Corning), washed, and replated overnight in 30μl AO medium as hanging drops. The next morning supernatants were collected and selected cytokines measured using an in-house developed and validated multiplex immunoassay (Laboratory of Translational Immunology, UMC Utrecht) based on Luminex technology (xMAP, Luminex Austin TX USA). The assay was performed as described previously^17^. Acquisition was performed with the Biorad FlexMAP3D (Biorad laboratories, Hercules USA) in combination with xPONENT software version 4.2 (Luminex). Data was analyzed by 5-parametric curve fitting using Bio-Plex Manager software, version 6.1.1 (Biorad).

### Neutrophil isolation and co-culture with AOs

Human neutrophils were isolated from sodium-heparin anticoagulated venous blood of healthy donors by density gradient centrifugation with and Ficoll (Amersham Biosciences). Erythrocytes were lysed in ammonium chloride buffer (155 mM NH4Cl; 10 mM KHCO3; 0.1 mM EDTA in double-distilled H2O; pH = 7.2) and neutrophils were resuspended in RPMI 1640 (Life Technologies, Paisley, UK) supplemented with 10% (v/v) HI FBS (Biowest, Nuaillé, France) and 50 U/ml Penicillin-Streptomycin (Thermo Fisher Scientific, Waltham, MA, USA). Purity of isolated neutrophils was analyzed using the CELL-DYN Emerald (Abbott Diagnostics, Illinois, USA). 100,000 freshly isolated neutrophils were co-seeded in BME with AOs that had been infected with/without RSV 3d earlier. Immediately after BME solidification, drops were overlaid with AO medium and live imaged as described. For quantification, neutrophils were labeled with 1μM Hoechst33342 prior to imaging with 3D confocal microscopy and the number of neutrophils attached to AOs with/without RSV counted 2h and 6h post cell-organoid-mixing.

### Generation of inducible dTomato and RSV-NS2/dTomato overexpressing AO lines

The following sequence comprising humanized NS1, GS linker, FLAG tag, HA tag, P2A sequence, humanized NS2, GS linker, V5 tag, 6xHis tag, and T2A sequence flanked by EcoRI restriction sites was synthesized by Integrated DNA Technologies (Leuven, Belgium) and cloned into IDT Vector Amp Blunt:

gaattcgccaccATGGGCAGCAACTCCCTGAGCATGATCAAAGTGCGGCTGCAGAACCTGTTCGACAACGACGAAGTCGCACTGCTCAAGATCACATGCTACACCGACAAGCTCATCCACCTGACCAACGCCCTGGCAAAGGCAGTGATCCACACTATCAAACTGAACGGTATCGTGTTCGTGCACGTCATCACCAGCAGCGACATCTGCCCTAACAACAACATCGTCGTCAAGTCCAACTTCACAACAATGCCCGTGCTGCAGAACGGCGGCTACATCTGGGAGATGATGGAGCTCACACACTGCTCCCAGCCCAACGGACTGATCGACGACAACTGCGAGATCAAGTTCTCCAAGAAGCTGAGCGACTCCACCATGACCAACTACATGAACCAGCTCTCCGAGCTGCTGGGATTCGACCTCAACCCTggcggcggcggctcgggaggaggaggaagcgactacaaggacgacgatgacaagTACCCCTACGACGTGCCCGACTACGCCggaagcggaGCTACTAACTTCAGCCTGCTGAAGCAGGCTGGAGACGTGGAGGAGAACCCTGGACCTATGGACACCACCCACAACGACACCACTCCACAGCGGCTGATGATCACCGACATGCGGCCACTGTCCCTGGAGACCACCATCACCTCCCTGACCCGCGACATCATCACCCACCGGTTCATCTACCTGATCAACCACGAGTGCATCGTGCGGAAGCTGGACGAGCGGCAGGCCACCTTCACCTTCCTGGTGAACTACGAGATGAAGCTGCTGCACAAAGTCGGCAGCACCAAGTACAAGAAGTACACTGAGTACAACACCAAATACGGCACCTTCCCAATGCCCATCTTCATCAACCACGACGGCTTCCTGGAGTGCATCGGCATCAAGCCCACAAAGCACACTCCCATCATCTACAAATACGACCTCAACCCTggtggtggtggttcaggaggaggatcgGGTAAGCCTATCCCTAACCCTCTCCTCGGTCTCGATTCTACGCATCATCACCATCACCACggaagcggaGAGGGCAGAGGAAGTCTGCTAACATGCGGTGACGTCGAGGAGAATCCTGGACCTgaattc. Humanized RSV-NS sequences were from Lo et al.^18^, 2A sequences from Kim et al.^19^. EcoRI-tdTomato-NotI was amplified from pCSCMV:tdTomato using primers aagaattcatggtgagcaagggcgagg and ctgcggccgcttacttgtacagctcgtccat and cloned into pLVX-TetOne^™^-Puro (Clontech) yielding an inducible dTomato lentiviral overexpression plasmid. EcoRI-hNS2-GS-V5-His-T2A-EcoRI was amplified from the IDT Vector using primers taccctcgtaaagaattcgccaccATGGACACCACCCACAACG and cttgctcaccatgaattcAGGTCCAGGATTCT and cloned into pLVX-TetOne-Puro-tdTomato yielding an inducible hNS2/dTomato lentiviral overexpression plasmid. Virus was generated and organoids infected as previously described^20^. Infected organoids were selected with 1μg⋅ml^-1^ puromycin (Sigma). Expression was induced with 5μM doxycycline (Sigma).

## Acknowledgements

We would like to acknowledge Carmen Lopez-Iglesias for supporting high pressure freezing and freeze substitution. We thank Raimond Ravelli for providing transmission EM overview software and Antoni P.A. Hendrickx for help with scanning EM. We acknowledge Anko de Graaff and the Hubrecht Imaging Center for supporting light microscopy. We thank Inez Bronsveld for obtaining intestinal biopsies. We thank the WKZ Pediatric Pulmonology Department for obtaining lavage fluid. We are grateful to Stieneke van der Brink and Nilofar Ehsani for preparing media. Genetically modified RSV variants were generously provided by Peter Collins, Mark E. Peeples, and Ulla Buchholz. We thank the Multiplex Core facility of the LTI UMC Utrecht for developing, validating, and performing immunoassays.

Work at the Hubrecht Institute was supported by MKMD and VENI grants from the Dutch Organization of Scientific Research NWO (ZonMw 114021012 and 916.15.182), a Stand Up to Cancer International Translational Cancer Research Grant (a program of the Entertainment Industry Foundation administered by the AACR) and an Alpe d’Huze/KWF program grant (2014-7006). The spinning disk microscope is funded by NWO equipment grant 834.11.002. Work in the laboratory of S.J.T. is part of the research program of the Foundation for Fundamental Research on Matter (FOM), as part of NWO. The work of G.H. and J.S.Z. was supported by a NWO VIDI grant (680-47-529) and a HFSP Career Development Award (CDA00076/2014-C). Work at the St. Antonius Hospital was supported by the Prof. Dr. Jaap Swierenga Foundation (development of human lung organoids). J.M.B., D.Z., and C.K.E. were supported by a grant from The Netherlands Organisation for Health Research and Development (ZonMW 40-00812-98-14103) and the Dutch Cystic Fibrosis Society (HIT-CF program).

## Author contributions

N.S. designed, performed, and analysed experiments and wrote the manuscript. N.S. and D.D.Z. designed, performed, and analysed swelling experiments. A.P. generated and analysed *TP53* mutant organoids. N.S., I.H., A.L., J.L., and A.O. performed and analysed RNA sequencing experiments. L.B. performed and analysed viability assays. N.S., D.K., F.W., M.F.M.O., K.K.D., E.F.S., E.E.V., C.H.M.M., C.K.E., S.F.B., R.G.V., and J.M.B. organized lung tissue collection. G.H.P., E.P.O, S.J.T., and J.S.Z. tracked and modelled single cell migration in RSV-infected organoids. N.S., L.T., and S.D. performed RSV infection experiments. J.K. and H.B. performed histology. M.F.M.O. classified lung tumors and evaluated organoid histology. K.K. and S.D. produced virus stocks. N.I. and P.J.P. performed transmission electron microscopy. M.C.V. quantified RSV replication. N.S. and E.R. generated organoid lines. M.L. isolated neutrophils. H.C. supervised the study.

## Conflict of interest

N.S., H.C., J.M.B. and C.K.E. are inventors on patents/patent applications related to organoid technology.

---

**Extended Data Figure 1.**
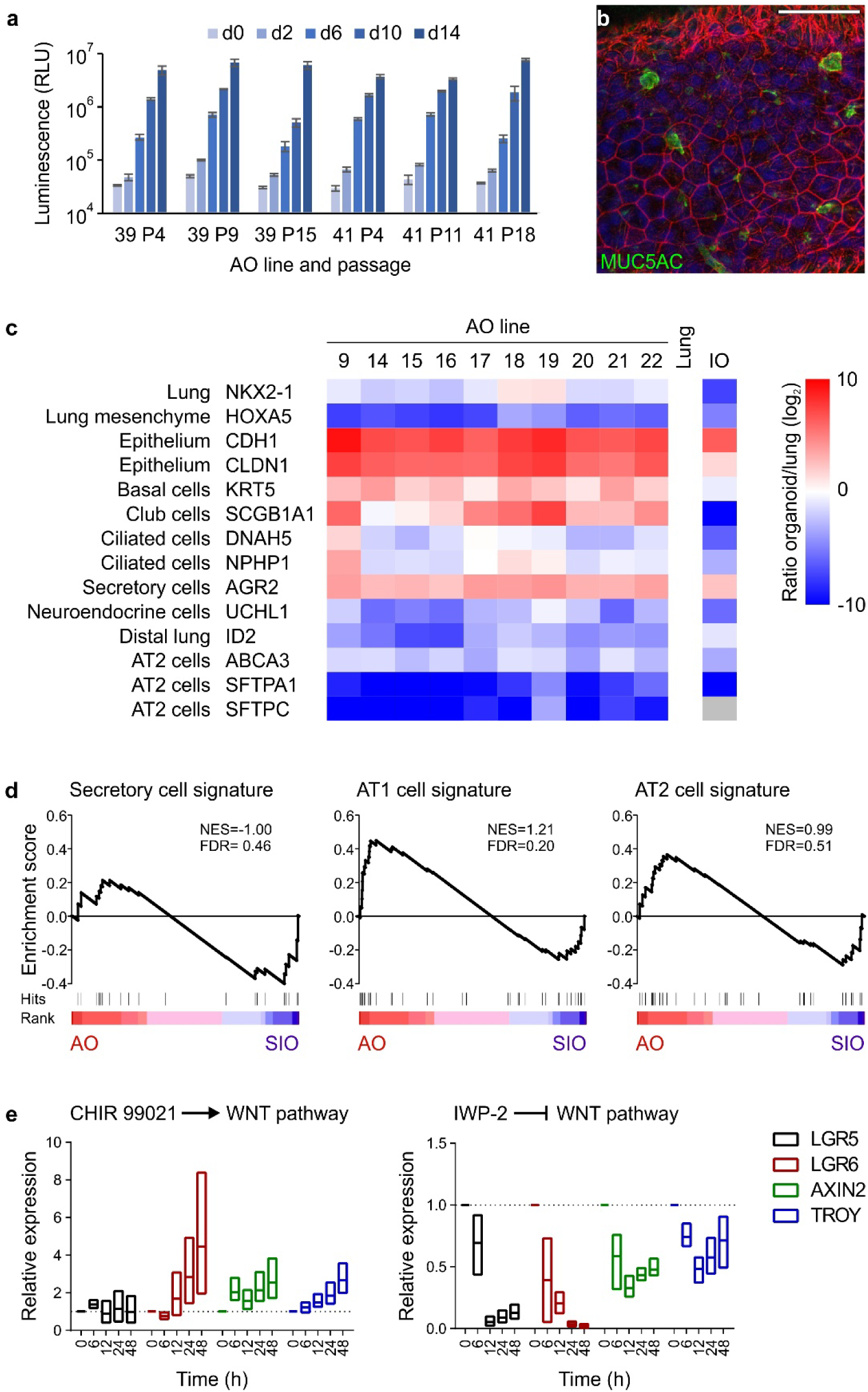
Further characterization of airway organoids. **a**, Luminescent cell viability assay comparing proliferative capacity of two independently generated AO lines at early, mid, and late passage numbers. Per group, 3000 cells were seeded and their expansion was measured at the indicated time points. Error bars represent standard deviations of technical triplicates. **b,** Immunofluorescent bottom-section of an AO showing secretory cell marker MUC5AC (green). Counterstained are the actin cytoskeleton (red) and nuclei (blue). Scale bar equals 50 μm. **c,** Heat map showing qPCR results of marker gene expression in ten individual AO lines compared to whole lung and one intestinal organoid line. Data are shown as log_2_- transformed ratio of sample over whole lung. AOs express lung cell marker NKX2-1 at comparable levels to whole lung (pale blue to pale red), while being negative for lung mesenchymal marker gene HOXA5 (dark blue) and strongly positive for general epithelial markers CDH1 and CLDN1 (dark red). AOs furthermore express relatively more KRT5 (basal cell marker), SCGB1A1 (club cell marker), and AGR2 (secretory cell marker). Ciliated cell markers DNAH5 and NPHP1 are expressed at similar levels in AOs and whole lung (pale blue to pale red). AOs express less of neuroendocrine marker UCHL1, distal lung marker ID2, and AT2 markers ABCA3, SFTPA1, and SFTPC (white to dark blue). The intestinal organoid line is positive for epithelial, basal, and secretory cell markers (pale red to red), and negative for general lung, lung mesenchyme, club cell, ciliated cell, neuroendocrine cell, distal lung, and AT2 cell markers (pale blue to dark blue). **d,** Gene set enrichment analysis plots showing no enrichment of the indicated gene signatures in transcriptomes of three AO lines compared to three small intestinal organoid (SIO) lines. NES = normalized enrichment score, FDR = false discovery rate. See auxiliary data table S1 for signatures and leading-edge genes. **e,** Box plots showing changes in the expression of stem cell marker and WNT target genes over time following manipulation of the WNT pathway. Upon strong activation of the WNT pathway using GSK-3 inhibitor CHIR 99021, LGR5 expression remains unaltered, while expression of AXIN2 and TROY is moderately and expression of LGR6 is strongly increased. Upon blocking the WNT pathway with porcupine inhibitor IWP-2, LGR5 and LGR6 expression drops sharply, while AXIN2 and TROY expression is decreased moderately. Analysis is based on three independent experiments using AO lines 15, 17, and 26.

**Extended Data Figure 2.**
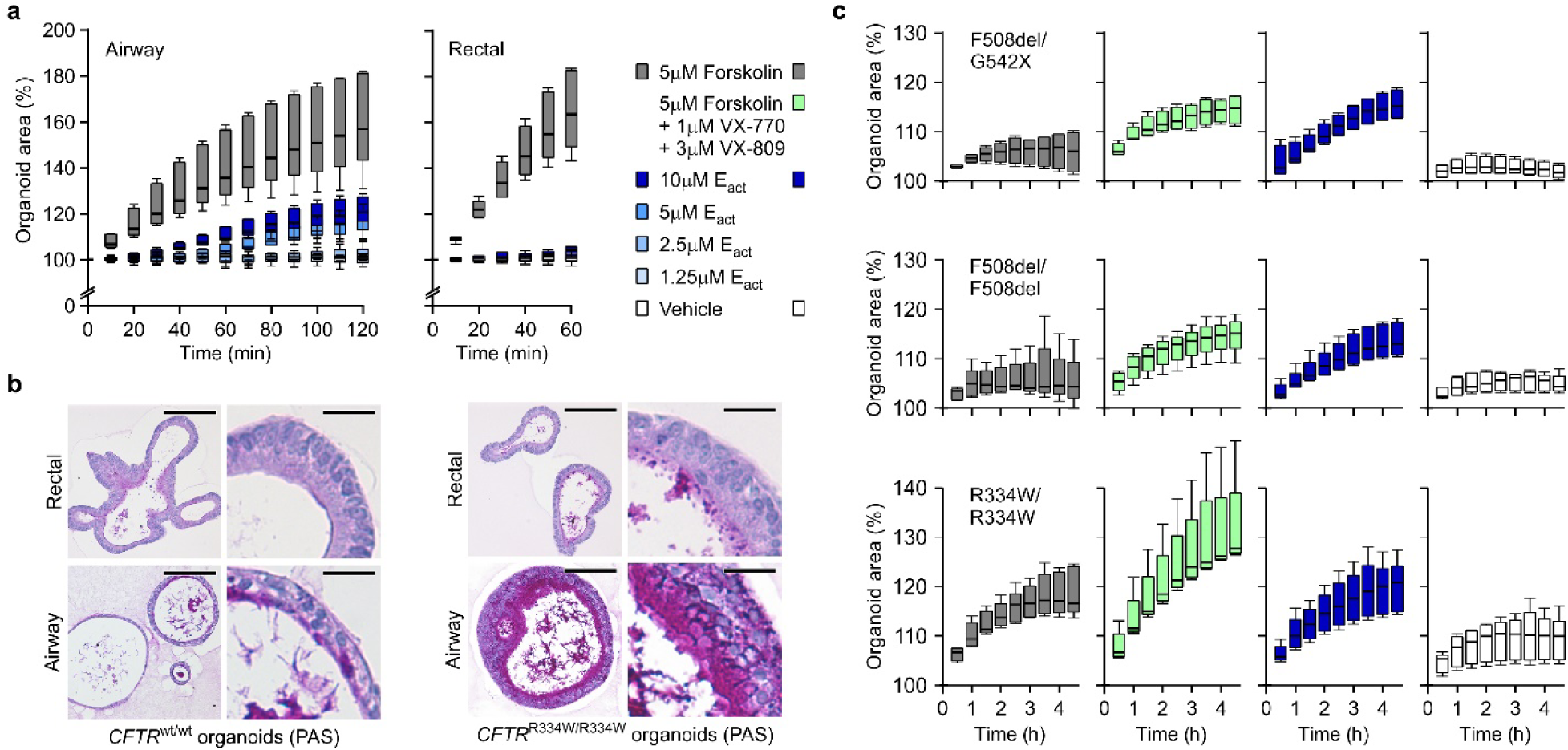
Additional data of airway organoids to study cystic fibrosis. **a**, Box-and-Whisker plots showing organoid swelling over time following stimulation with forskolin or E_act_. While forskolin causes swelling in both organoid types (grey boxes), E_act_ causes concentration-dependent swelling only of AOs (blue hued boxes). Shown are pooled data from three different AO and two different rectal organoid lines used in three to four independent experiments. See Figure 2a, b for the respective AUC plots. **b**, Representative histological sections of PAS-stained wild-type organoids (unmatched, left panel) and organoids from a CF patient with *CFTR*^R334W/R334W^ mutation (right panel). PAS-positive mucus is occasionally present within wild-type AOs but not rectal wild-type organoids, while the CF patient AOs regularly show thick layers of PAS-positive mucus. Rectal organoids from the same CF patient display only occasional regions with PAS-positive mucus. Rectal organoids were generated from rectal biopsies, AOs were generated from lung resection (wild-type) or BAL-fluid (CF patient). Scale bars equal 50 μm (overviews) and 10 μm (details). See Figure 2c for PAS-stained *CFTR*^F508del/F508del^ organoid sections. **c**, Box-and-Whisker plots showing CF patient AO swelling over time following addition of the indicated stimuli. The respective CFTR mutations are given atop every row of plots. Forskolin-induced swelling (grey boxes) does not exceed vehicle controls in AOs with *CFTR*^F508del/G542X^ and *CFTR*^F508del/F508del^ genotypes, but increases in the presence of VX-770 and VX-809 (green boxes). In the same organoids, E_act_-induced swelling (blue boxes) exceeds forskolin-induced swelling to a similar extend. AOs with the milder *CFTR*^R334W/R334W^ genotype (bottom row) show moderate forskolin-induced swelling that is increased in the presence of VX-770 and VX-809 and paralleled by E_act_-induced swelling. Shown are pooled data of four to five independent experiments. See Figure 2d for the corresponding AUC-plots.

**Extended Data Figure 3.**
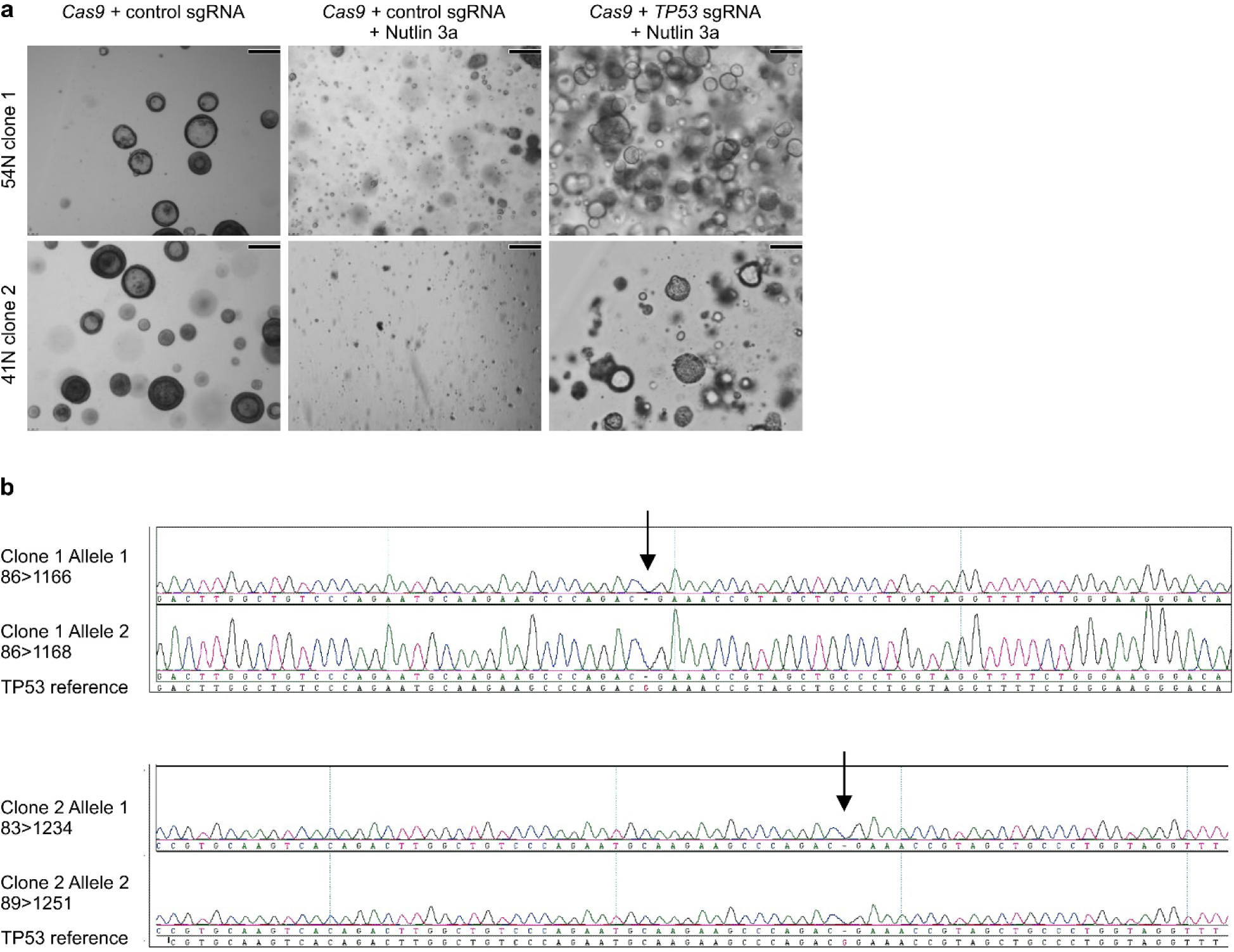
Additional data of *TP53* edited AOs. **a**, Bright field images showing the effect of Nutlin 3a selection on two independently *TP53* gene edited AO clones. While control AOs do not expand under selection (middle column), *TP53* edited AO clones do (right column). Scale bars equal 100 μm. **b**, Sequencing chromatograms of *TP53* edited AO clones showing bi-allelic generation of *TP53* frame shifts.

**Extended Data Figure 4.**
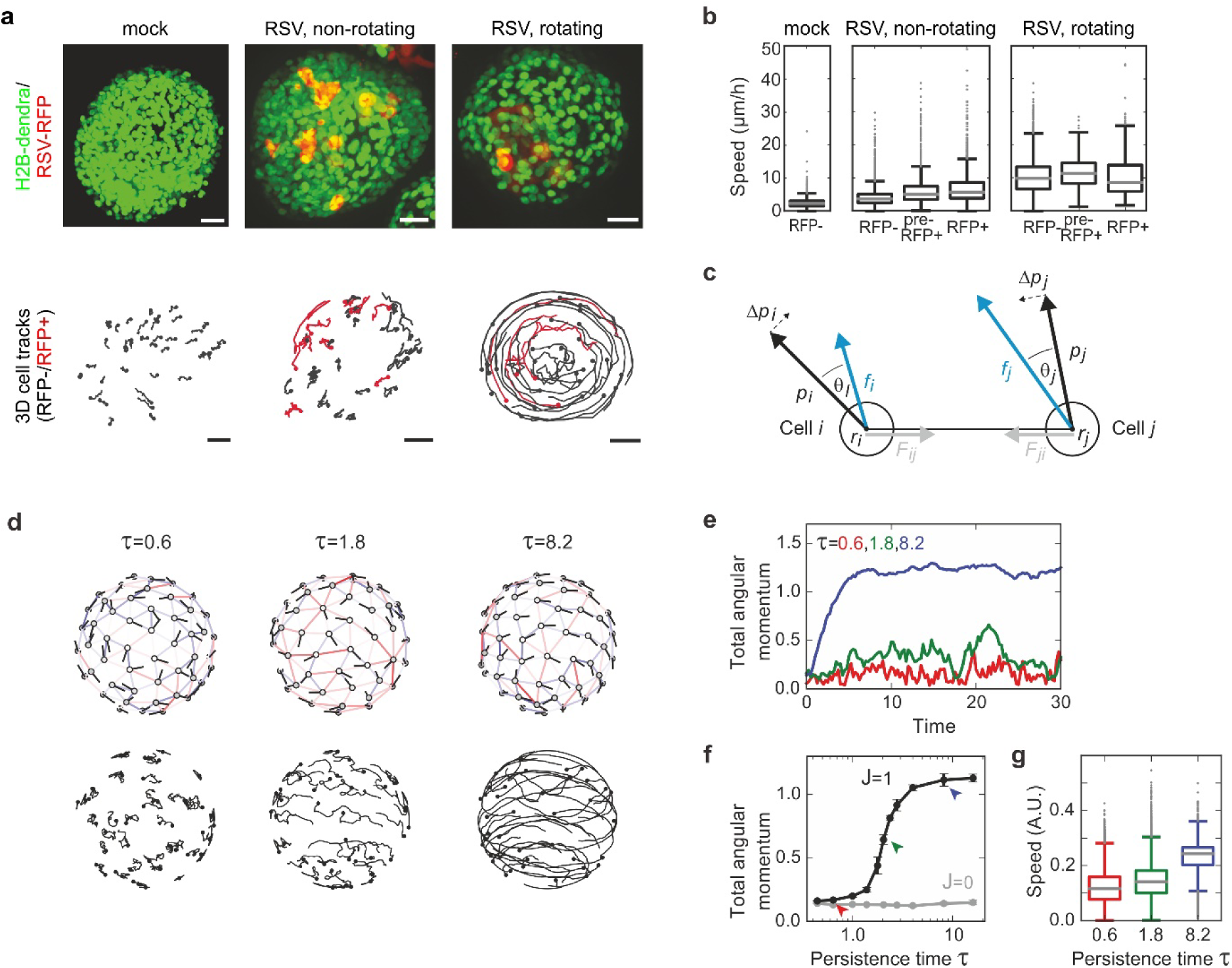
Single cell velocities of RSV-infected AOs and mathematical modeling of organoid rotation. **a**, Examples of non-infected (left), RSV-infected and non-rotating (middle), and RSV-infected and rotating (right) organoids with the corresponding tracks of randomly selected nuclei (n=41, 33, 35 respectively). Track durations are 14 h. Circles indicate the starting position of each track, and line color stands for either RPF- (black) or RFP+ (red) cells. Scale bars represent 25μm. **b**, Box plots of the speed distribution of every tracked nucleus at each time point for non-infected (n=3), infected but non-rotating (n=6) and infected and rotating (n=3) organoids, where 15-45 individual nuclei were tracked per organoid. For the infected organoids, nuclei were classified as RFP-when they showed no RFP signal for the duration of the track, pre-RFP+ when the nuclei showed no RFP signal yet but became RFP+ later, and RFP+ for nuclei that showed RFP signal. **c**, Vector schematic depicting the modeled relation between cells migrating within the constraints of an organoid sphere. See methods for details. **d**, Snapshots of the cell configuration (left) and cell tracks (right) for simulations with n=100 cells and increasing persistence time τ=0.6, 1.8 and 8.2. The persistence time indicates the mean time over which the cell maintains its direction of polarization, in the absence of cell-cell interactions. White markers represent cell centers, with black lines showing the direction of the polarization vector. Adjacent cells are connected by springs, which are shown as colored lines. Springs are red when stretched and blue when compressed. Cell tracks are shown for the same time period for all three simulations, with black circles indicating the starting position of each track. See also Supplementary Video 7. **e**, Total angular momentum of the cell configuration as a function of time for simulations starting with random initial distribution of polarity vectors. For sufficiently high persistence time (**τ**=8.2, blue line), the cells rapidly establish rotational motion. **f**, Total steady state angular momentum as function of the persistence time, for simulations with cell-cell communication (black, *J*=1) and without (grey, ***J***=0). Colored arrows indicate the persistence times corresponding to the simulations in panel e. **g**, Box plots of the distribution of cell speed for the different persistence times in panel e. The distribution is calculated from n=5 independent simulations.

**Extended Data Figure 5.**
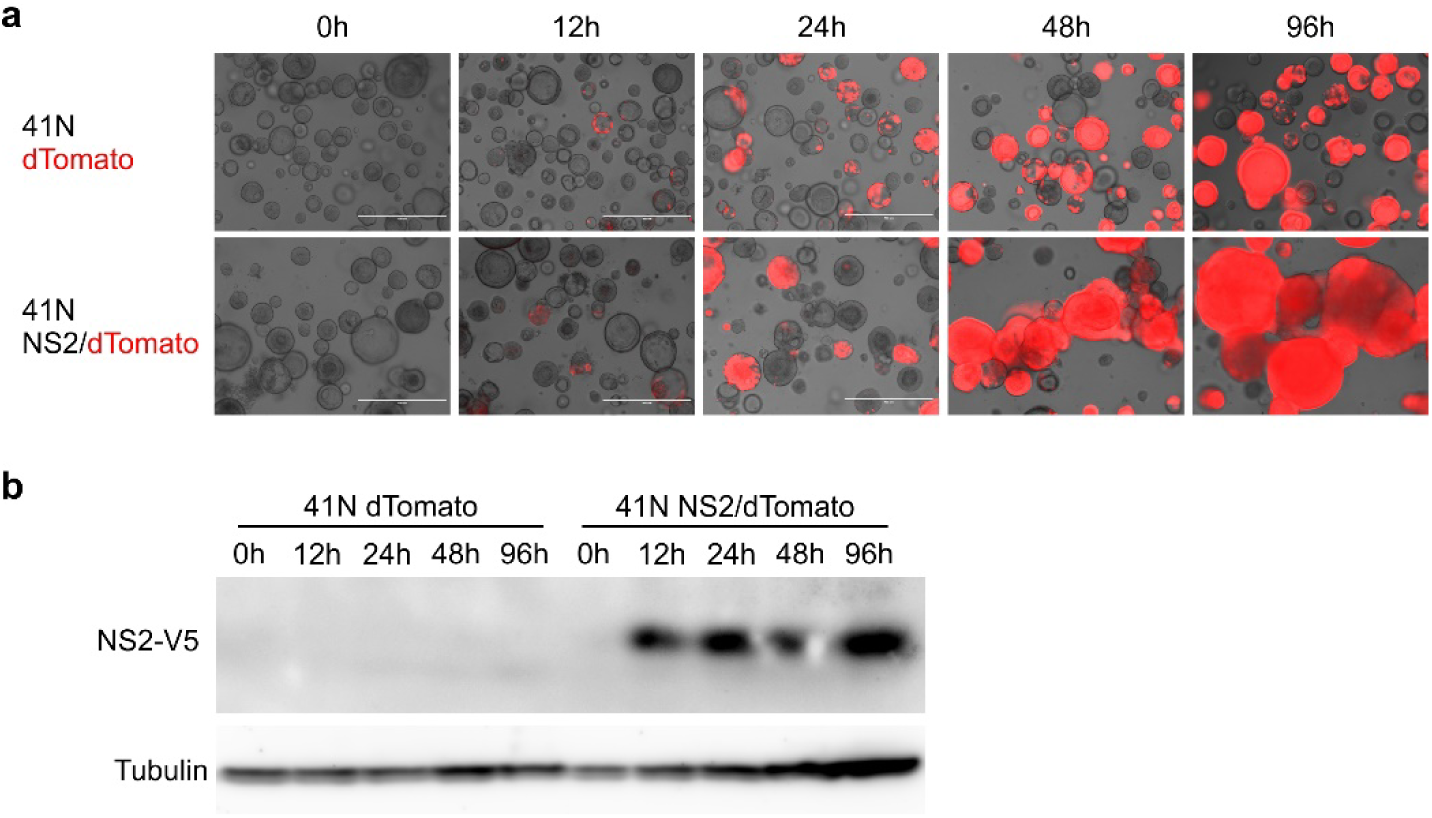
Inducible expression of NS2 in AOs. **a**, Brightfield/fluorescent micrographs of AOs inducibly overexpressing dTomato (top) or NS2 and dTomato (bottom) taken at the indicated time points following stimulation with doxycycline. While red signal increases in both lines, organoid fusion exclusively takes place in AOs expressing NS2. See Supplementary Video 11. Scale bars equal 400 μm. **b**, Western blots of protein lysates from the indicated AOs taken at the indicated time points following stimulation with doxycycline. NS2 protein is robustly detectable after 12h of stimulation.

## Supplementary Tables

Supplementary Table 1 – Airway organoid media recipe

Supplementary Table 2 – Transcriptome analysis AO vs SIO.

Supplementary Table 3 – Hotspot cancer gene sequencing of tumoroids

Supplementary Table 4 – Transcriptome analysis mock vs RSV-infected AOs.

## Supplementary Videos

Supplementary Video 1 – Sequentially taken transmission micrographs through an AO.

Supplementary Video 2 – Time-lapse bright field microscopy of ciliar movement within an AO.

Supplementary Video 3 – Time-lapse bright field microscopy of whirling mucus within an AO.

Supplementary Video 4 – Time-lapse bright field and fluorescence microscopy of AOs following infection with RSV-GFP (green). Note increasing GFP signal over time, high organoid motility, and fusion events in RSV-GFP infected AOs.

Supplementary Video 5 – Time-lapse confocal microscopy of AOs (H2B-dendra, green) following mock infection. AOs expand over time due to proliferation (note fast resolving mitotic figures) but otherwise remain static in place.

Supplementary Video 6 – Time-lapse confocal microscopy of AOs (H2B-dendra, green) following infection with RSV-RFP (red). Note increasing RFP signal over time, high cell motilities, organoid rotation, and loss of epithelial cohesion in RSV-RFP infected AOs.

Supplementary Video 7 – 3D-simulation of the effect of increased persistence times (τ) of cell motilities on organoid rotation. Low and medium persistence times do not result in organoid rotation (left and middle, τ=0.6 and τ=1.8 respectively), whereas high persistence time (right, τ=8.2) does.

Supplementary Video 8 – Time-lapse bright field and fluorescence microscopy of Mock-infected AO (top) vs RSV-RFP-infected AO (bottom) in the presence of primary human neutrophils. Left bright field, right RFP-channel. Note the increased migration of neutrophils towards the RSV-infected organoid.

Supplementary Video 9 – Time-lapse confocal microscopy of Mock-infected AO (left, light green, spherical nuclei) vs RSV-RFP-infected AO (right, bright red, light green) in the presence of primary human neutrophils (bright green, lobulated nuclei). Bright white indicates apoptotic cells (SYTOX). Note the increased number of neutrophils present on the RSV-infected organoid.

Supplementary Video 10 – Time-lapse bright field and fluorescence microscopy of AOs micro-injected with wild-type RSV-GFP (top) or mutant RSV-GFP lacking NS2 (bottom). Note the lack of both cell motility and GFP signal-increase in the ΔNS2 condition.

Supplementary Video 11 – Time-lapse bright field and fluorescence microscopy of AOs inducibly overexpressing dTomato (top) or NS2 together with dTomato (bottom). While dTomato signal increases in both AO lines, organoid rotation is exclusively observed in NS2 overexpressing AOs.

